# A self-organized signaling hierarchy patterns the cortex during cell repair

**DOI:** 10.64898/2026.08.21.746176

**Authors:** Lila E Hoachlander-Hobby, Alison Moe, Tim Liang, Yuming Liu, Adriana E Golding, Kathryn P McCauley, Trang T Pham, Tom Burke, Peter Bieling, Kevin W Eliceiri, Matthew E Larson, William M Bement

## Abstract

Cells generate dynamic patterns of Rho GTPase activation to direct the subsequent patterning of Rho GTPase effectors needed to remodel the cell cortex during processes ranging from cell division to cell repair. To understand how such patterns arise, we used live cell imaging, time-resolved Rho GTPase manipulations, and a novel computational tool to study the spatiotemporal dynamics of Rho, Cdc42 and several downstream Cdc42 targets in wounded *Xenopus laevis* oocytes. We find that the characteristic wound-induced segregation of Rho and Cdc42 activity into concentric zones is followed by polarization of the Cdc42 zone such that Toca-1 progressively concentrates at the back of the Cdc42 zone while Arp2/3, cofilin, cortactin, and the Rho GAP p190RhoGAP progressively concentrate at the front of the Cdc42 zone, where it overlaps the Rho zone. Remarkably, the juxtaposition of Rho activity to Cdc42 is required for the polarization of p190RhoGAP, while p190RhoGAP is responsible for establishing the boundary between the Cdc42 and Rho zones. The results indicate that the characteristic segregation of the Rho and Cdc42 zones, as well as the polarization of the Cdc42 zone, arise from cortical self-organization. Further, these findings reveal a simple mechanism for hierarchical establishment of cortical patterns: recruitment of new proteins to regions of signaling compartment overlap.

## Introduction

Cells routinely experience damage to their plasma membrane and the underlying cortical cytoskeleton. Such damage may occur in the course of their natural function [1], as a result of physical trauma [2], or due to pathological agents and conditions [3]. Accordingly, cells possess conserved repair mechanisms in which an inrush of calcium through the damage site initiates resealing of the plasma membrane [4,5] and restoration of the cortical cytoskeleton [6,7]. A growing body of evidence indicates that these processes are medically important, in that deficits in cell repair result in or exacerbate several major diseases including muscular dystrophies [8–10] and diabetes [11].

In addition to its medical relevance, cell repair has emerged as a powerful model to study polarization and patterning of the cell cortex, as damage triggers rapid accumulation of signaling and cytoskeletal proteins [12–14] and lipids [15,16] around wound sites. One of the fundamental polarization events elicited by cell damage is local activation of Rho GTPases. In budding yeast [17], Dictyostelium [18], C. elegans epidermis [19], Drosophila syncytial embryos [20], Xenopus oocytes and embryos [12,21], and cultured human muscle cells [22], either Rho, Cdc42, Rac or all three are focally activated at cellular wounds. One of the fundamental cortical patterning events elicited by cell damage is formation of concentric rings (often called “zones”) of polarized proteins and lipids.

In wounded mouse myofibrils, F-actin concentrates in a ring slightly displaced from the wound edge, which circumscribes a region rich in calcium-dependent membrane binding proteins such as annexins and dysferlin [23]; in wounded Drosophila syncytial embryos, F-actin and the Rho GTPases Rac and Cdc42 concentrate in a ring slightly displaced from the wound edge while Rho is concentrated in an overlapping ring that is positioned closer to the wound edge [13,20]; in wounded Xenopus oocytes [12], F-actin and Cdc42 activity are concentrated in a ring slightly displaced from the wound edge while Rho activity is concentrated in a ring at the immediate wound edge. Exactly how such concentric organization arises is a matter of intense interest, both because it informs our understanding of cortical pattern formation and because manipulations that disrupt the patterns impair cell healing [20,24].

In a previous study of the contributions of crosstalk to the positioning and length scales of the Rho and Cdc42 activity zones, we found that the level of active, GTP-bound Rho (hereinafter Rho-T) controls the position of the Cdc42 zone while the level of active, GTP-bound Cdc42 (hereinafter Cdc42-T) controls the width of the Rho zone [24]. The latter finding was particularly intriguing, in that while it was clear that Cdc42-T acted upstream of something (presumably a Rho GAP) that limits the spread of the Rho zone away from the wound edge, the presumptive site of Rho inactivation did not precisely track with Cdc42-T, but instead, was positioned slightly in front (i.e. closer to the wound edge) of the Cdc42-T peak.

Here, we have developed an automated system for spatiotemporal analysis of wounds and used it to characterize the dynamics of Cdc42-T targets to understand the mechanism of Cdc42-T-mediated Rho-T confinement. We find that wound patterning is far more intricate than previously realized, with the Cdc42-T zone developing a distinct front-back polarity within ∼90s of wounding, such that different targets sort to the front or back of the zone. We further show that one of the front-polarized downstream targets of Cdc42-T is p190RhoGAP, which forms an enzymatic corral that prevents the spread of Rho-T away from the wound edge. Finally, we show that polarization of p190RhoGAP is dependent on Rho-T itself, indicating that the patterning of the array is self-organized.

## Results

### A computational tool for objective, automated analysis of spatiotemporal protein dynamics during cell repair

Protein and phospholipid dynamics during single cell repair have previously been quantified by radially averaging signal intensity around wounds over time [24,25]. However, this method is time-intensive, difficult to render unbiased, and generally limits the amount of information obtained. Inspired by previous efforts to automate quantification of fluorescent signal during multicellular wound repair [26,27], we developed a novel computational tool that permits automated spatial averaging of signal intensity at different distances from the wound edge over time. This tool, named the Sphinctalyzer, uses a single reference frame of a time-lapse movie and a user-drawn reference path around any zone of enriched protein to average signal intensities of pixels equidistant from the reference ring through the entire time-lapse movie (Figs 1A and S1A). This approach, based in part on the Live-wire algorithm and the snakes-based MEDUSA tool [27,28], outputs line-scans and spatiotemporal data for each frame of the movie and can be used to batch-process multiple movies in a row. To make this tool broadly useful, we also developed a graphical user interface (GUI), which simply requires the investigator to import one or more movies, adjust parameters for each movie, and run the automated analysis. The Sphinctalyzer GUI is user-friendly, supports multi-channel movie analysis, allows for parameters to be updated for each movie, displays line-scans and spatiotemporal results, and has various output and post-processing options to meet analysis needs.

**Fig 1.**
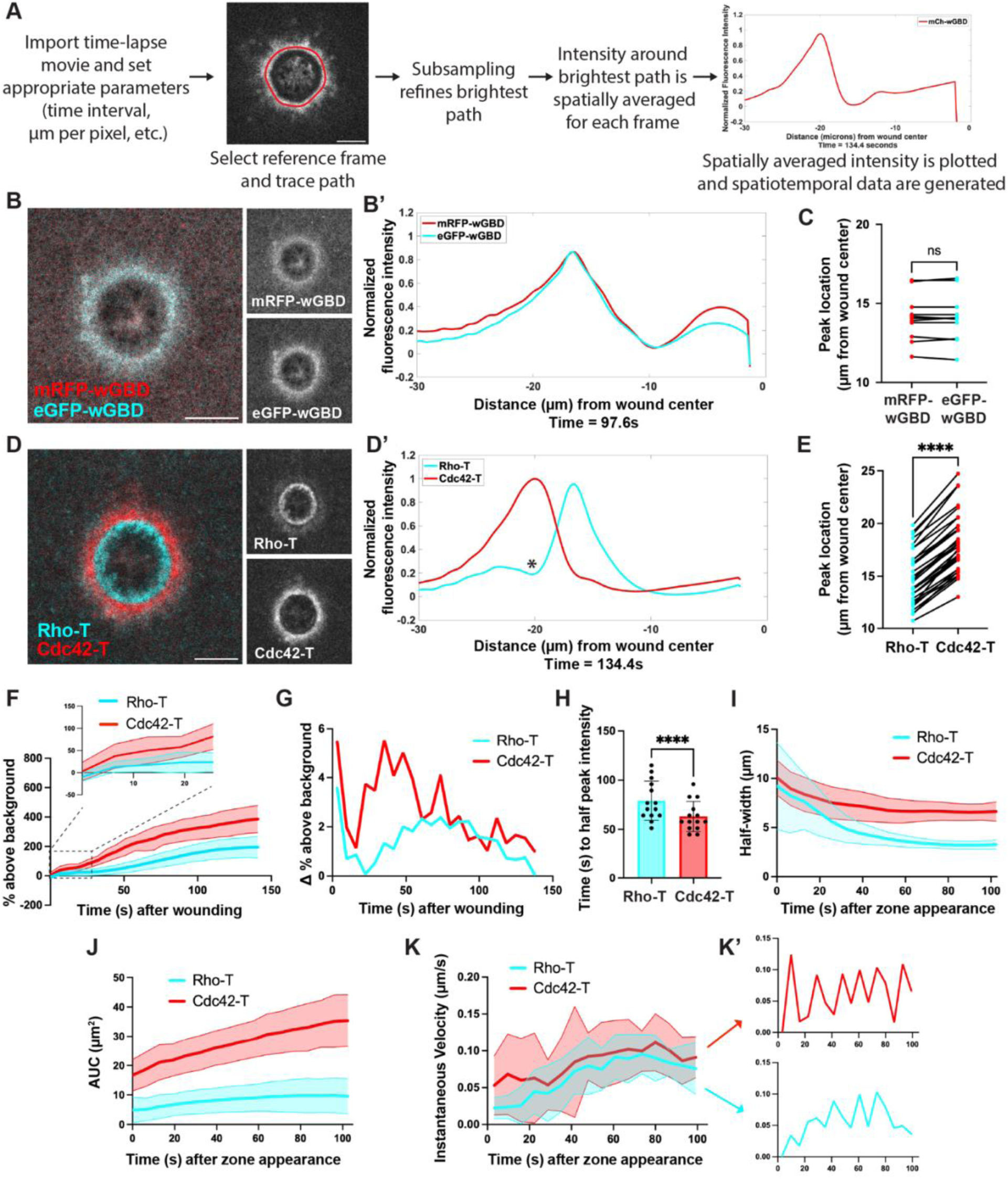
Spatial averaging reveals spatiotemporal dynamics of Rho GTPases during cell repair. **(A)** Brief overview of Sphinctalyzer workflow and data output. **(B)** Two probes for Cdc42-T, mRFP-wGBD and eGFP-wGBD, colocalize around wounded *Xenopus laevis* oocytes. **(B’)** Spatially averaged and normalized signal intensity of mRFP-wGBD and eGFP-wGBD from the micrograph in (B). Wound center is 0 on the x-axis. **(C)** Peak locations of the mRFP-wGBD and eGFP-wGBD zones relative to the wound center at 25% wound closure (n=11). **(D)** Probes for Rho-T and Cdc42-T, dTom-rGBD and eGFP-wGBD respectively, enrich in concentric zones around wounded *Xenopus laevis* oocytes. **(D’)** Spatially averaged and normalized signal intensity of Rho-T and Cdc42-T from the micrograph in (D). Wound center is 0 on the x-axis. The asterisk denotes the dip in Rho-T activity. **(E)** Rho-T and Cdc42-T zone peak locations relative to the wound center at 25% wound closure (n=33). **(F)** Rho-T and Cdc42-T zone intensity measured as % above background over time after wounding (n=14). Inset shows Cdc42-T intensity rising faster than Rho-T intensity immediately after wounding. **(G)** First derivative of data from (F) to show the rate of change in Rho-T and Cdc42-T intensity over time after wounding. **(H)** Rho-T and Cdc42-T zone time to half peak intensity (n=14). **(I)** Rho-T and Cdc42-T zone width measured as full width at half maximum (half-width) over time after zone appearance (n=14). **(J)** Rho-T and Cdc42-T zone area measured as area under the curve (AUC) over time after zone appearance (n=14). **(K)** Rho-T and Cdc42-T zone instantaneous velocities measured as change in zone position from one time point to the next over time after zone appearance. (n=14). **(K’)** Representative data from one cell used in K. Graphs have been split to show Cdc42-T (red, top) and Rho-T (cyan, bottom) instantaneous velocities (same axes as in K) separately. Note the oscillatory changes in instantaneous velocity. (H) shows the mean +/− SD and paired t-tests were used in C, E, and H. In (F), (I), (J), and (K) the thick line is the average, and the shaded regions above and below are the SD. ****p<0.0001. Scale bars for micrographs = 20µm.

To assess the fidelity of data extracted by the Sphinctalyzer from time-lapse movies, we first tested its output for two proteins expected to localize to the same position around wounds: mRFP-wGBD and eGFP-wGBD, two Cdc42 activity probes that bind to Cdc42-T [12] and only differ in the fluorophore used for visualization. Dynamic colocalization of mRFP-wGBD and eGFP-wGBD is qualitatively apparent in a representative micrograph (Figs 1B, S1B and S1B’). The Sphinctalyzer faithfully reports that mRFP-wGBD and eGFP-wGBD localize to the same position around wounds as shown by an individual line-scan (Fig 1B’) and by quantification of multiple experiments (Fig 1C).

As an additional test, we used the Sphinctalyzer to compare the dynamics of Cdc42-T (detected with eGFP-wGBD) to Rho-T (detected with dTom-rGBD), as Cdc42-T and Rho-T form concentric rings around wounds, with Cdc42-T circumscribing Rho-T [12,24] (Fig 1D). Consistent with previous findings, the Sphinctalyzer-generated line-scan shows two spatially distinct activity zones (Fig 1D’) with the Cdc42-T zone significantly farther from the wound center than the Rho-T zone is (Fig 1E). However, the Sphinctalyzer also permits automated quantification of important features that were previously difficult to assess. For example, normalizing the fluorescence intensity (to the peak intensity achieved during repair) makes direct visual comparison of different peak positions simpler, particularly in cases where the proteins differ in the degree to which they are concentrated within zones (e.g. Figs 1D’ and S1C). Moreover, the normalization also revealed a dip in Rho activity in the Cdc42-T zone that was not previously detectable (denoted by asterisks in Figs 1D’ and S1C). Upon inspection, ∼72% of cells had a Rho-T dip (59/82 cells), which appeared an average of 83 seconds after wounding (83.13 +/−22.14s); the significance of this observation will be apparent below. In addition, the Sphinctalyzer output demonstrated that the Cdc42-T rise is initially much sharper than that of the Rho-T, as shown by the % above background over time (Fig 1F), the change in % above background over time (Fig 1G), and the time to half peak intensity (Fig 1H). Moreover, the Sphinctalyzer revealed that both the Cdc42-T and the Rho-T zones start out at around 10 µm in width but then narrow to ∼8 and 4 µm, respectively, within 1 minute of their first appearance (Fig 1I). Comparison of these results to quantification of the area under the curve of the zones (Fig 1J) showed that the while the Rho-T zone increased modestly in total Rho activity even as it is narrowing, the Cdc42-T zone increased more dramatically in total Cdc42 activity as it narrowed.

Lastly, the Rho-T and Cdc42-T zone instantaneous velocity was investigated to better understand Rho GTPase zone translocation throughout repair. The instantaneous velocity represents the change in Rho-T and Cdc42-T zone position from one time point to the next and is displayed over time to graphically visualize zone acceleration. The Rho-T and Cdc42-T instantaneous velocities and overall acceleration were similar when quantified from multiple experiments, suggesting that the concentric zones move in a coordinated fashion throughout repair (Fig 1K). Surprisingly, however, inspection of data from individual experiments revealed oscillatory changes in both Rho-T and Cdc42-T instantaneous velocities over time (Rho-T period (s): 17.85 +/− 6.25; Cdc42-T period (s): 17.43 +/− 4.91) (Figs 1K’ and S1D). This suggests that either the actomyosin contraction that powers wound closure [29] or the waves of Rho and Cdc42 activity [30], or both are inherently oscillatory, a point addressed below. Collectively, the above data show that the Sphinctalyzer can recapitulate prior findings and reveal previously unknown features of Rho GTPase dynamics.

### Cdc42-T defines the Rho-T zone boundary, width, and total activity, while Rho-T controls the Cdc42-T zone translocation

Previous work showed that fusion proteins based on the calcium- and phospholipid-binding C2 domain of protein kinase C β (hereinafter “C2”) are rapidly recruited from the cytosol to wound edges [31]. Because C2 is soluble prior to wounding, this approach permits disruption of particular targets only where and when cells are wounded. This avoids disruption of the cortical cytoskeleton prior to wounding that can arise from chronic suppression of the Rho GTPases by other approaches, such as expression of dominant negative Rho GTPase mutants or depletion of Rho GTPases or their GEFs. To exploit the Sphinctalyzer for analysis of Rho GTPase crosstalk, we first monitored Rho activity following expression of a Cdc42 GAP catalytic domain (Chn1GAP) fused to C2 (hereinafter “Chn1GAP-C2”). Consistent with previous results [24], this manipulation significantly reduced Cdc42 activity (Figs 2A and S2A). The Sphinctalyer revealed that Cdc42-T suppression with Chn1GAP-C2 had no significant effect on the rate of wound closure, nor on the oscillation period of the Rho-T zone (control period (s): 16.29 +/− 5.33; Chn1GAP-C2 period (s): 16.98 +/− 4.41) (Fig 2B) but it did result in expansion of Rho activity away from the wound edge (Figs 2A, 2A’, S2B, and S2C)[24].

**Fig 2.**
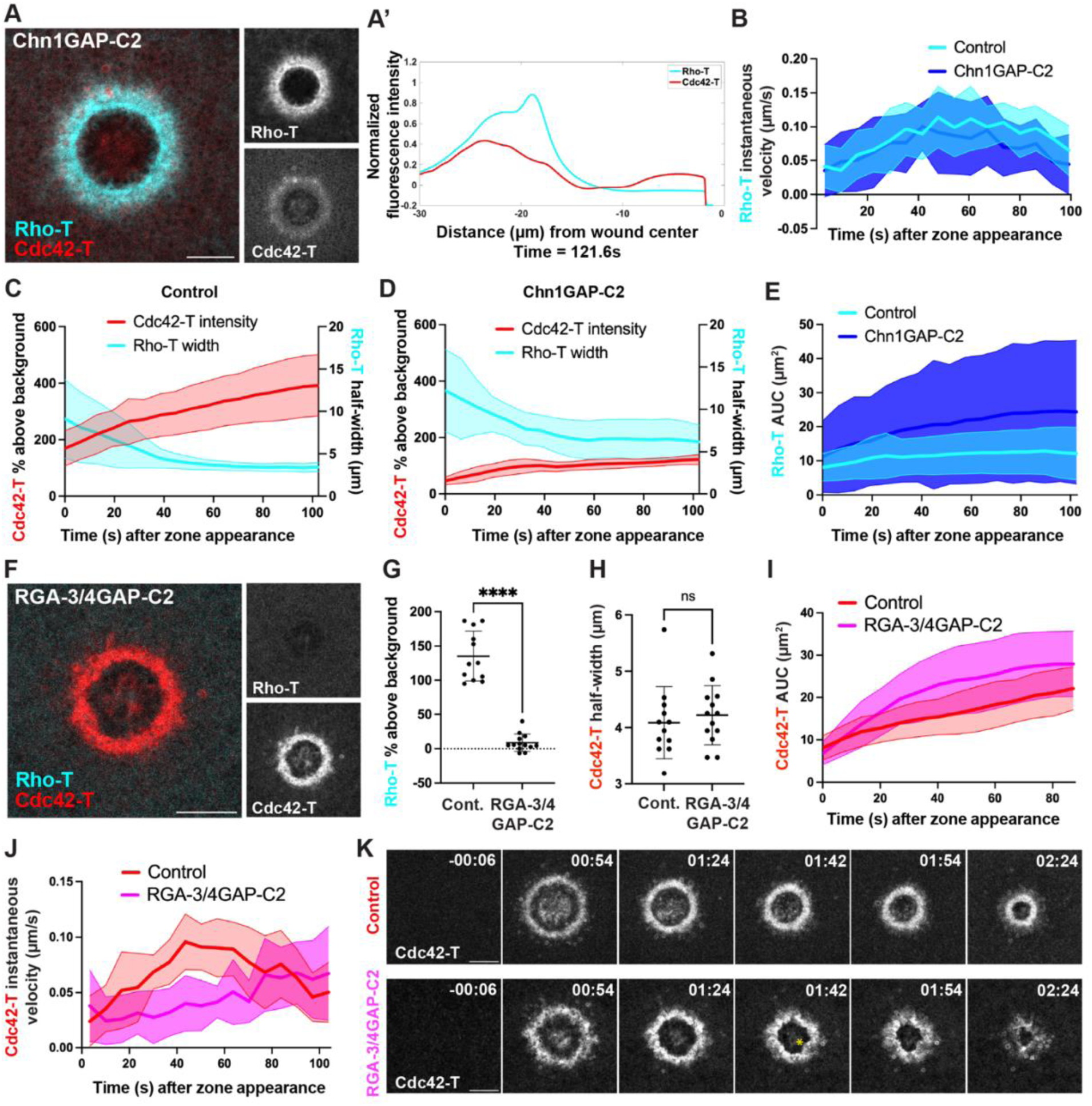
Cdc42 activity establishes Rho-T zone width, boundary, and total activity and Rho activity mediates Cdc42-T zone translocation. **(A)** Chn1GAP-C2 expression diminishes Cdc42-T and results in spread of Rho activity away from the wound edge. **(A’)** Spatially averaged and normalized signal intensity of Rho-T and Cdc42-T from the micrograph in (A). Wound center is 0 on the x-axis. **(B)** Rho-T zone instantaneous velocity over time after zone appearance in controls (n=14) vs. cells expressing Chn1GAP-C2 (n=8). **(C)** Rho-T zone width compared to Cdc42-T zone intensity over time after zone appearance in controls (n=14). **(D)** Rho-T zone width compared to Cdc42-T zone intensity over time after zone appearance in cells expressing Chn1GAP-C2 (n=8). **(E)** Rho-T zone area in controls (n=14) vs. cells expressing Chn1GAP-C2 (n=8). **(F)** RGA-3/4GAP-C2 expression diminishes Rho-T but does not affect the Cdc42-T zone boundary. **(G)** Rho-T zone intensity at 25% wound closure in controls (n=12) vs. cells expressing RGA-3/4GAP-C2 (n=13). **(H)** Cdc42-T zone width at 20% wound closure in controls (n=12) vs. cells expressing RGA-3/4GAP-C2 (n=13). **(I)** Cdc42-T zone area over time after zone appearance in controls (n=12) vs. cells expressing RGA-3/4GAP-C2 (n=13). **(J)** Cdc42-T zone instantaneous velocity over time after zone appearance in controls (n=12) vs. cells expressing RGA-3/4GAP-C2 (n=13). **(K)** Montage of Cdc42-T zone in controls (top row) vs. cells expressing RGA-3/4GAP-C2 (bottom row). Note that the inner edge of the Cdc42-T zone is more circular in controls, whereas the inner edge has an irregular shape in cells expressing RGA-3/4GAP-C2 (denoted by asterisk). Time is in minutes:seconds. (G) and (H) show the mean +/− SD and unpaired t-tests were used. In (B), (C), (D), (E), (I), and (J) the thick line is the average, and the shaded regions above and below are the SD. ****p<0.0001. Scale bars for micrographs = 20µm.

To better understand how Cdc42 activity affects the Rho-T zone boundary, Cdc42-T zone intensity and Rho-T zone width were compared over time. In control cells, there was an inverse relationship between these features: as Cdc42-T intensity rises, the Rho-T zone narrows (Figs 2C and S2D (r = −0.5520)). However, in cells expressing Chn1GAP-C2, Cdc42-T intensity remained low and there was a significant increase in Rho-T zone width throughout repair relative to control cells (Figs 2D and S2C). Further, there was also a higher average Rho-T zone area under the curve relative to controls that was evident immediately upon the onset of Rho-T zone formation (Fig 2E), showing that reducing Cdc42 activity leads to more total Rho activity around the wound. These findings indicate that Cdc42 activity normally suppresses Rho activity shortly after the onset of Rho-T zone formation, early in the repair process. They also indicate that the entire region around the wound can support Rho activity when Cdc42-T is suppressed.

To permit reciprocal experiments involving time-resolved Rho-T suppression, we generated a wound-targeted Rho GAP based on the catalytic domain of the Rho GAP RGA-3/4 [32] fused to C2 (hereinafter “RGA-3/4GAP-C2”). RGA-3/4GAP-C2 significantly reduced Rho activity around the wound (Fig 2F and 2G), however, this did not affect Cdc42-T zone width (Fig 2H). Suppressing Rho activity did however increase Cdc42-T zone area under the curve and intensity over time, beginning at about 10s after initial Cdc42-T zone formation and about 1 minute after wounding, respectively (Figs 2I, S2E, and S2F). This finding was missed in previous studies wherein this metric could not be assessed [12,24]. Together, these data suggest that the Cdc42-T zone, in contrast to the Rho-T zone, is self-limiting in terms of width and that Rho activity suppresses Cdc42 activity not by limiting the spread of the Cdc42-T zone, but rather by limiting Cdc42 activity within the Cdc42-T zone itself.

To better understand Cdc42-T zone translocation in the absence of Rho-T-stimulated actomyosin contraction, Cdc42-T zone instantaneous velocity over time was compared in controls and cells expressing RGA-3/4GAP-C2. RGA-3/4GAP-C2 expression led to slower Cdc42-T zone instantaneous velocity early in repair (i.e. before 75 seconds after zone appearance) (Fig 2J), suggesting that Rho-T-mediated actomyosin contraction is important for Cdc42-T zone acceleration. This did not, however, affect the oscillatory changes in Cdc42-T zone instantaneous velocity (S2G Fig) (control period (s): 17.53 +/− 4.66; RGA-3/4GAP-C2 period (s): 17.45 +/− 5.09), which implies that the oscillations are intrinsic to the Rho GTPase waves, rather than actomyosin contraction per se. Consistent with previous results obtained via injection of C3 exotransferase [30], most wounds still closed following RGA-3/4GAP-C2 expression, albeit in an irregular, crawling-like fashion (Fig 2K and S1 Video).

### Direct Cdc42-T targets localize to the Cdc42-T zone

Previous work in wounded oocytes demonstrated that Cdc42-T is excluded from the Rho- T zone by the Rho-T-dependent recruitment of Abr, which inactivates Cdc42-T via its GAP domain [33]. However, the means by which Cdc42-T suppresses Rho activity is unknown, in part because the signal transduction pathway downstream of Cdc42-T has not been characterized in this system. As a first step to address this point, the spatiotemporal dynamics of three direct targets (effectors) of Cdc42-T were investigated: Neural Wiskott-Aldrich Syndrome Protein (N-WASP), Transducer of Cdc42-dependent actin assembly-1 (Toca-1), and p21 activated kinase 2 (Pak2). N-WASP concentrated at wounds in the middle of the Cdc42-T zone (Figs 3A, 3B, S3A, and S3B) on average within 0.03 +/− 0.20µm from the Cdc42-T zone peak, and reached half peak intensity significantly later than Cdc42-T (Figs 3C and 3D). Additionally, Chn1GAP-C2 expression significantly reduced N-WASP recruitment around wounds (S3M Fig), as expected if N-WASP is recruited to wounds in a Cdc42-T-dependent manner.

**Fig 3.**
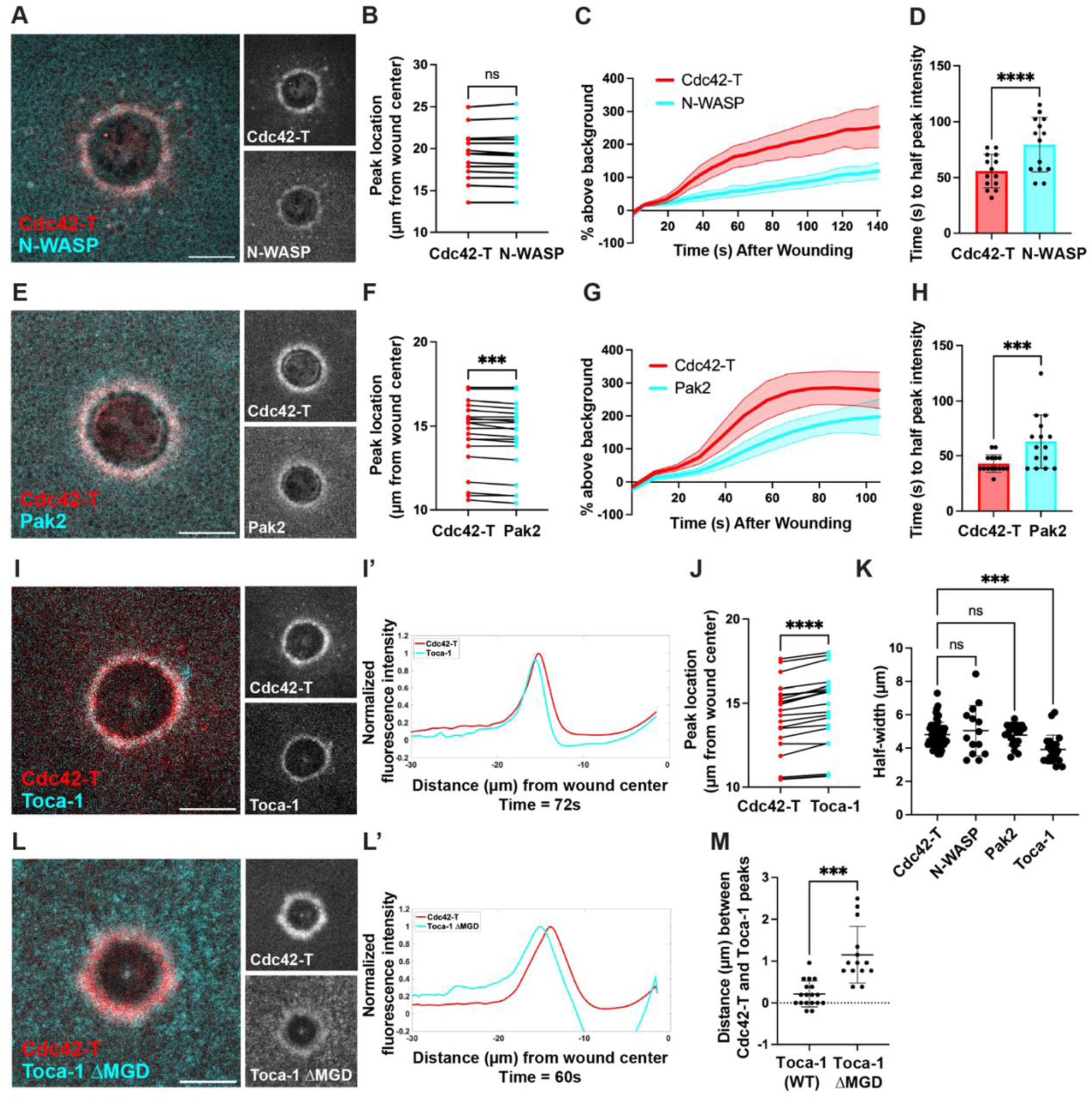
Spatiotemporal dynamics of direct Cdc42-T targets during repair. **(A)** N-WASP enriches around wounds in the Cdc42-T zone. **(B)** Cdc42-T and N-WASP peak location at 25% wound closure (n=14). **(C)** Cdc42-T and N-WASP intensity over time after wounding (n=14). **(D)** Cdc42-T and N-WASP time to half peak intensity (n=14). **(E)** Pak2 enriches around wounds in the Cdc42-T zone. **(F)** Cdc42-T and Pak2 peak location at 25% wound closure (n=20). **(G)** Cdc42-T and Pak2 intensity over time after wounding (n=15). **(H)** Cdc42-T and Pak2 time to half peak intensity (n=15). **(I)** Toca-1 enriches around wounds in the Cdc42-T zone. **(I’)** Spatially averaged and normalized signal intensity of Cdc42-T and Toca-1 from the micrograph in (I). Wound center is 0 on the x-axis. **(J)** Cdc42-T and Toca-1 peak location at 25% wound closure (n=21). **(K)** Cdc42-T (n=55) vs. N-WASP (n=14), Pak2 (n=20), and Toca-1 (n=21) widths at 25% wound closure. **(L)** Toca-1ΔMGD localizes farther outside the Cdc42-T zone than WT Toca-1 does. **(L’)** Spatially averaged and normalized signal intensity of Cdc42-T and Toca-1ΔMGD from the micrograph in (L). Wound center is 0 on the x-axis. **(M)** Distance between the Cdc42-T and WT Toca-1 peaks (n=18) vs. the distance between the Cdc42- T and Toca-1ΔMGD peaks (n=14) at 25% wound closure. (D), (H), (K), and (M) show the mean +/− SD. In (C) and (G) the thick line is the average, and the shaded regions above and below are the SD. Paired t-tests were used in (B), (D), (F), (H), and (J). One-way ANOVA with a Dunnett comparison was used in (K). An unpaired t-test was used in (M). ***p<0.001, ****p<0.0001. Scale bars for micrographs = 20µm.

Pak2 also concentrated at wounds in the Cdc42-T zone (Fig 3E) with a slight bias toward the front (leading edge) of the zone (Figs 3F, S3D, and S3E), as the Pak2 peak was on average shifted 0.12 +/− 0.12µm closer to the wound center than the Cdc42-T zone peak. Like N-WASP, Pak2 reached half peak intensity significantly later than Cdc42-T (Fig 3G and 3H). Both N-WASP and Pak2 half-width narrowed over time (S3C and S3F Fig) and did not differ from Cdc42-T zone half-width (Fig 3K).

Toca-1 also concentrated at wounds in the Cdc42-T zone (Fig 3I), but in contrast to N-WASP and Pak2, it consistently occupied a narrower region at the back (trailing edge) of the Cdc42-T zone (Figs 3I’, 3J, 3K, S3G, and S2 Video), with the Toca-1 peak on average shifted 0.43 +/− 0.24µm farther from the wound center than the Cdc42-T zone peak. A similar rearward shift was also evident when Toca-1 was compared to Pak2 instead of Cdc42-T (S3H and S3H’ Fig), indicating that this pattern was not a consequence of using the Cdc42-T probe as a marker. Toca-1 reached half peak intensity significantly later than Cdc42-T (S3I and S3J Fig).

Since wounding elicits the formation of a ring-like microdomain of phosphatidylinositol 4,5-bisphosphate (PIP2) that circumscribes and partially overlaps the Cdc42-T zone [15], and Toca-1 binds PIP2 [34,35], we reasoned that Toca-1 may be targeted to the back of the Cdc42-T zone via overlapping input from both PIP2 and Cdc42-T. To test this notion, we compared the localization of Cdc42-T to Toca-1 ΔMGD, a mutant with reduced affinity for Cdc42-T [36]. Consistent with this idea, Toca-1 ΔMGD still concentrated around wounds (Fig 3L), but its peak activity shifted significantly backward (i.e. farther from the wound center) compared to WT Toca-1, such that the peak of Toca-1 ΔMGD localized slightly outside the Cdc42-T zone (Figs 3L’, 3M, and S3K). Toca-1 ΔMGD half-width also increased in comparison to WT Toca-1 (S3L Fig).

### Arp2/3, cofilin, and cortactin concentrate at the front of the Cdc42- T zone

To gain further insight into Cdc42-T signaling during the wound response, we next characterized the spatiotemporal dynamics of several common downstream Cdc42-T targets: the Actin-related protein (Arp2/3) complex, which is activated by N-WASP [34,37–39]; cofilin, which promotes F-actin disassembly and is typically enriched in regions of branched actin networks [40]; and cortactin, which binds and stabilizes F-actin branches generated by Arp2/3 [41–43].

Consistent with these proteins serving as downstream Cdc42-T targets, Arp2/3, cofilin and cortactin all concentrated around the wound in the Cdc42-T zone (Fig 4A, 4E, and 4I) and reached half peak intensity significantly later than Cdc42-T (Figs 4B, 4F, 4J, S4A, S4D, and S4G). Moreover, their recruitment was dependent on Cdc42-T, as shown via expression of Chn1GAP-C2 (Fig 4M, 4N, and S4J). Throughout repair, the Arp2/3 and cortactin regions were consistently narrow (S4B and S4H Fig), whereas the cofilin zone began wide and narrowed over time (S4E Fig). Overall, the Arp2/3, cortactin, and cofilin zones were significantly narrower than the Cdc42-T zone (S4K Fig).

**Fig 4.**
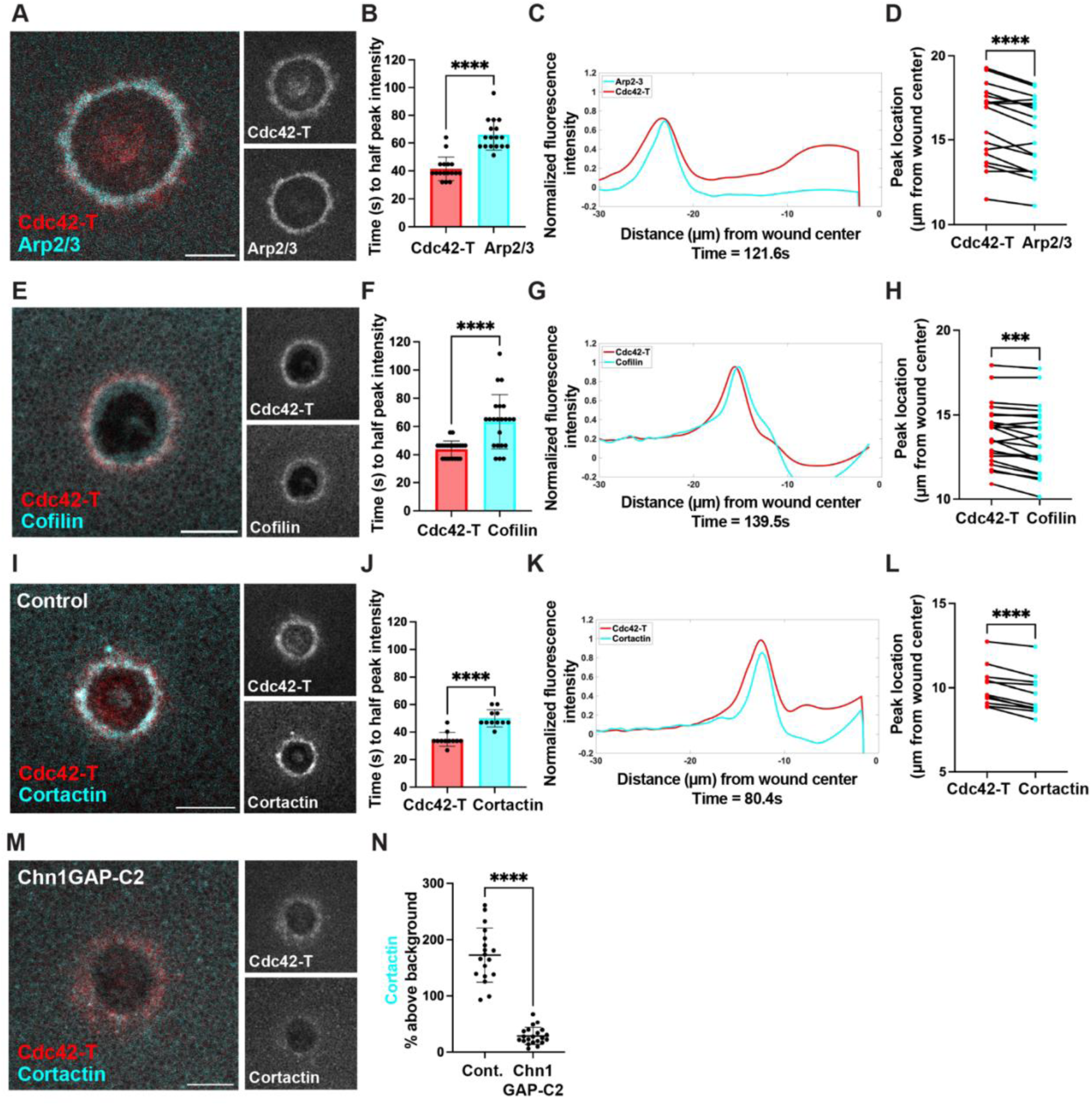
Indirect Cdc42-T targets polarize to the front of the Cdc42-T zone. **A)** Arp2/3 enriches around wounds in the Cdc42-T zone. **(B)** Cdc42-T and Arp2/3 time to half peak intensity (n=17). **(C)** Spatially averaged and normalized signal intensity of Cdc42-T and Arp2/3 from the micrograph in (A). Wound center is 0 on the x-axis. **(D)** Cdc42-T and Arp2/3 peak location at 25% wound closure (n=17). **(E)** Cofilin enriches around wounds in the Cdc42-T zone. **(F)** Cdc42-T and cofilin time to half peak intensity (n=22). **(G)** Spatially averaged and normalized signal intensity of Cdc42-T and cofilin from the micrograph in (E). Wound center is 0 on the x-axis. **(H)** Cdc42-T and cofilin peak location at 25% wound closure (n=22). **(I)** Cortactin enriches around wounds in the Cdc42-T zone. **(J)** Cdc42-T and cortactin time to half peak intensity (n=11). **(K)** Spatially averaged and normalized signal intensity of Cdc42-T and cortactin from the micrograph in (I). Wound center is 0 on the x-axis. **(L)** Cdc42-T and cortactin peak location at 25% wound closure (n=11). **(M)** Chn1GAP-C2 expression diminishes both Cdc42-T and cortactin. **(N)** Cortactin intensity at 25% wound closure in controls (n=17) vs. cells expressing Chn1GAP-C2 (n=21). (B), (F), (J), and (N) show the mean +/− SD. Paired t-tests were used in (B), (D), (F), (H), (J), and (L). An unpaired t-test was used in (N). ***p<0.001, ****p<0.0001. Scale bars for micrographs = 20µm.

Remarkably, Arp2/3, cofilin, and cortactin all progressively localized to the front (leading edge) of the Cdc42-T zone (S3, S4, and S5 Videos). That is, while initially appearing either in the center or slightly behind the center of the Cdc42-T zone, by 20-30 s after their appearance in distinct zones, Arp2/3, cofilin, and cortactin became concentrated at the front of the Cdc42-T zone (Figs 4C, 4D, 4G, 4H, 4K, 4L, S4C, S4F, and S4I). A similar frontward shift was also evident when cofilin was compared to Pak2 instead of Cdc42-T (S4L and S4L’ Fig), indicating that this pattern was not a consequence of using the Cdc42-T probe as a marker. Thus, just as the Toca-1 localization pattern suggests that the Cdc42-T zone develops a distinct back, the behavior of Arp2/3, cofilin, and cortactin indicate that it also develops a distinct front.

### The Rho-T zone is required for forward polarization of cortactin in the Cdc42-T zone

The results above indicated that the rearward polarization of Toca-1 within the Cdc42-T zone arises from the juxtaposition of the circular PIP2 microdomain with the Cdc42-T zone. We therefore sought to test the possibility that the forward polarization of downstream Cdc42-T targets such as cortactin arises from the juxtaposition of the Rho-T zone with the Cdc42-T zone. To test this possibility, we suppressed Rho activity via expression of RGA-3/4GAP-C2. In contrast to controls, wherein cortactin localized to a narrow region at the front of the Cdc42-T zone (Fig 4), in cells expressing RGA-3/4GAP-C2, cortactin remained spread throughout the Cdc42-T zone (Fig 5A, 5B and 5C), leading to a significant increase in cortactin half-width (Fig 5D). Suppression of Rho activity via microinjection of C3 exotransferase also suppressed polarization of cortactin (S5 Fig). Curiously, while disruption of the Cdc42-T-Rho-T juxtaposition had no effect on the Cdc42-T zone width (Fig 2H), the total amount of cortactin around wounds increased (Fig 5E and 5F). This suggests that some aspect of the polarization process selectively removes cortactin from the cortex.

**Fig 5.**
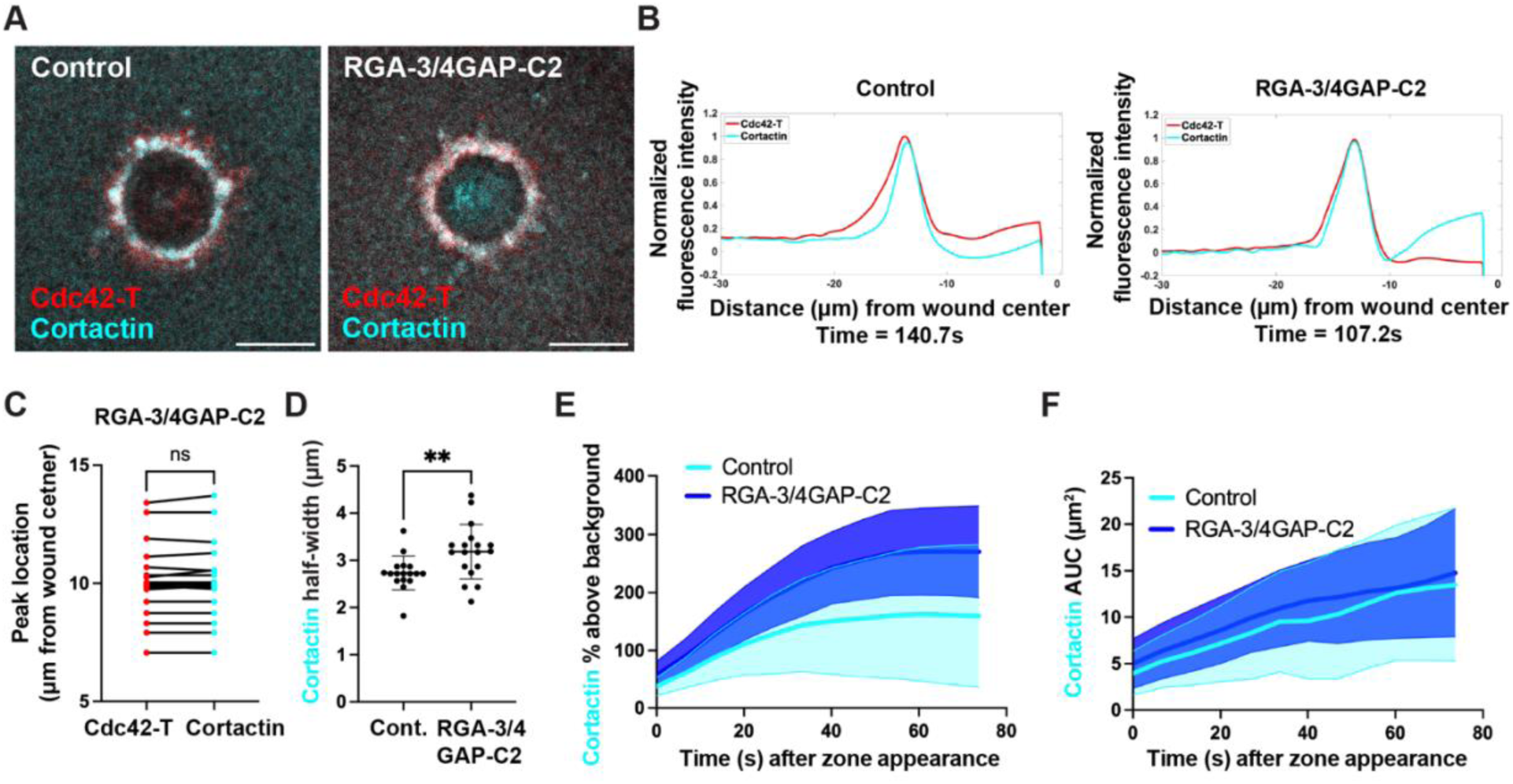
Cortactin polarization to the front of the Cdc42-T zone depends on Rho activity. **(A)** (Left) cortactin localizes to the front of the Cdc42-T zone in controls. (Right) cortactin localizes throughout the Cdc42-T zone when Rho-T is inhibited via RGA-3/4GAP-C2 expression. **(B)** (Left) spatially averaged and normalized signal intensity of Cdc42-T and cortactin from the control micrograph in (A). (Right) spatially averaged and normalized signal intensity of Cdc42-T and cortactin from the RGA-3/4GAP-C2 micrograph in (A). Wound center is 0 on the x-axes. **(C)** Cdc42-T and cortactin peak location at 20% wound closure in cells expressing RGA-3/4GAP-C2 (n=18). **(D)** Cortactin half-width at 20% wound closure in controls (n=17) vs. cells expressing RGA-3/4GAP-C2 (n=18). **(E)** Cortactin intensity over time after zone appearance in controls (n=14) vs. cells expressing RGA-3/4GAP-C2 (n=20). **(F)** Cortactin area over time after zone appearance in controls (n=14) vs. cells expressing RGA-3/4GAP-C2 (n=20). (D) shows the mean +/− SD. In (E), and (F), the thick line is the average, and the shaded regions above and below are the SD. A paired t-test was used in (C), and an unpaired t-test was used in (D). **p<0.01. Scale bars for micrographs = 20µm.

### p190RhoGAP regulates Rho-T zone confinement

The demonstration that Cdc42-T suppression results in spreading of Rho activity away from the wound (Fig 2) [24] indicates that Cdc42-T signaling somehow inactivates Rho-T, thereby confining Rho activity to the wound edge. Because the peak of Cdc42-T itself is usually positioned rearward to the Rho-T dip (see above), the impact of Cdc42-T on Rho-T is unlikely to be direct. However, the discovery of the frontward polarization of downstream Cdc42-T targets suggests the possibility that something associated with the leading edge of the Cdc42-T zone is responsible for Rho inactivation. Because cortactin was previously reported to interact with p190RhoGAP (hereinafter “p190”), a GAP that inactivates Rho-T [44], we tested the potential involvement of p190 in Rho-T patterning. P190 co-localized with cortactin around wounds (Figs 6A, 6B, and S6A) at the front of the Cdc42-T zone (S6B and S6C Fig and S6 Video) and arrived at the wound slightly after cortactin (Fig 6C and S6D), suggesting that cortactin might recruit p190 to the Cdc42-T zone. P190 circumscribed Rho-T (Figs 6D and S6E), and just as suppression of Cdc42-T with Chn1GAP-C2 suppressed cortactin recruitment (Fig 4M and 4N), it also suppressed p190 recruitment, leading to spread of Rho activity away from the wound edge (Fig 6E and 6F). Further, comparison of controls to samples in which Cdc42-T was suppressed showed a strong inverse correlation between the levels of p190 and the half-width of the Rho-T zone (Figs 6G (r = −0.5848), S6F, and S6G).

**Fig 6.**
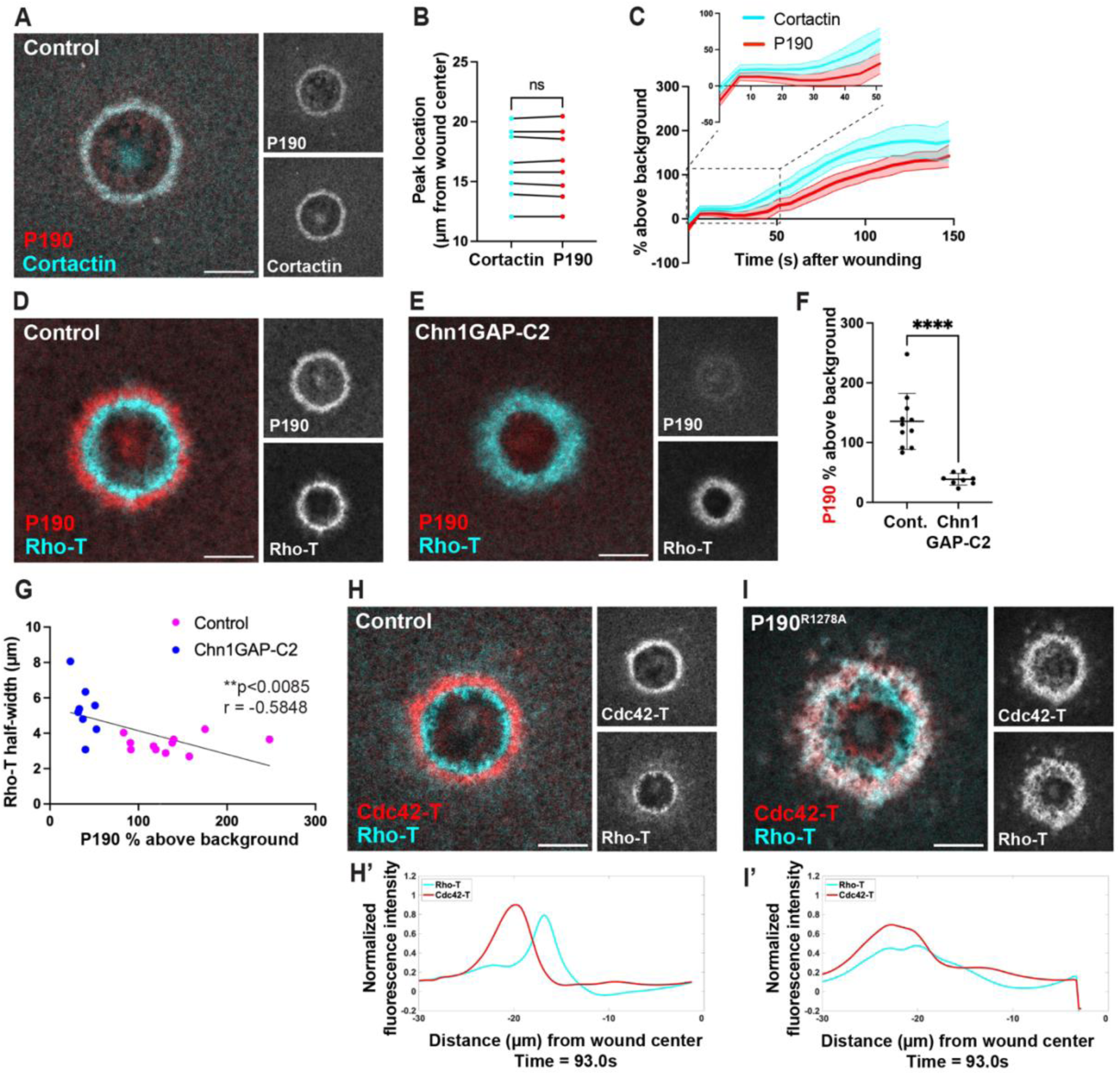
The cortactin-associated p190RhoGAP confines Rho-T to the wound edge. **(A)** p190RhoGAP (p190) enriches around wounds and co-localizes with cortactin. **(B)** Cortactin and p190 peak location at 25% wound closure (n=8). **(C)** Cortactin and p190 intensity over time after wounding (n=8). Inset shows cortactin intensity rising slightly earlier than p190 intensity. **(D)** p190 circumscribes the Rho-T zone. **(E)** Chn1GAP-C2 expression diminishes p190 and results in spread of Rho-T away from the wound edge. **(F)** P190 intensity at 25% wound closure in controls (n=11) vs. cells expressing Chn1GAP-C2 (n=8). **(G)** Correlation of Rho-T half-width and P190 intensity at 25% wound closure (n=19). **(H)** Cdc42-T and Rho-T form mutually exclusive concentric zones in controls. **H’)** Spatially averaged and normalized signal intensity of Rho-T and Cdc42-T from the micrograph in (H). Wound center is 0 on the x-axis. **(I)** P190^R1278A^ expression results in spread of Rho activity away from the wound edge and into the Cdc42-T zone. **(I’)** Spatially averaged and normalized signal intensity of Rho-T and Cdc42-T from the micrograph in (I). Wound center is 0 on the x-axis. (F) shows the mean +/− SD. In (C), the thick line is the average, and the shaded regions above and below are the SD. A paired t-test was used in (B) and an unpaired t-test was used in (F). Pearson correlation and simple linear regression were used in (G). ****p<0.0001. Scale bars for micrographs = 20µm.

As a direct test of whether p190 regulates Rho-T confinement, we generated a dominant negative p190 mutant (p190^R1278A^) that can bind Rho-T, but cannot inactivate Rho-T [45,46]. p190^R1278A^ expression resulted in dramatic spread of Rho-T away from the wound edge such that Rho activity infiltrated the Cdc42-T zone and was no longer confined to the immediate wound edge (Fig 6H-I’ and S7 and S8 Videos). This led to a significantly smaller separation between the Cdc42-T and Rho-T zones (S6H and S6I Fig), wider Rho-T zone (S6J and S6K Fig), and greater Rho-T zone area under the curve (S6L and S6M Fig). Cumulatively, these results show that p190 is a downstream target of Cdc42-T signaling that regulates Rho-T confinement by inactivating Rho-T in the Cdc42-T zone.

### Rho-T mediates polarization of its inhibitor, p190RhoGAP, at the front of the Cdc42-T zone

The localization pattern of p190 at the front of the Cdc42-T zone (S6B and S6C Fig), as well as its known interaction with cortactin suggested that it too might polarize in a Rho-T-dependent manner. To test this idea, we monitored p190 dynamics following Rho suppression via expression of RGA-3/4GAP-C2. As observed for cortactin, this manipulation abolished polarization of p190 (Fig 7A, 7B, and 7C). Surprisingly, unlike cortactin, Rho-T suppression resulted in a significantly smaller p190 half-width and slightly lower intensity and area under the curve over time compared to controls (Fig 7D, 7E, and 7F). These results show that Rho-T confinement ultimately results from Rho-T-dependent recruitment of its own inhibitor.

**Fig 7.**
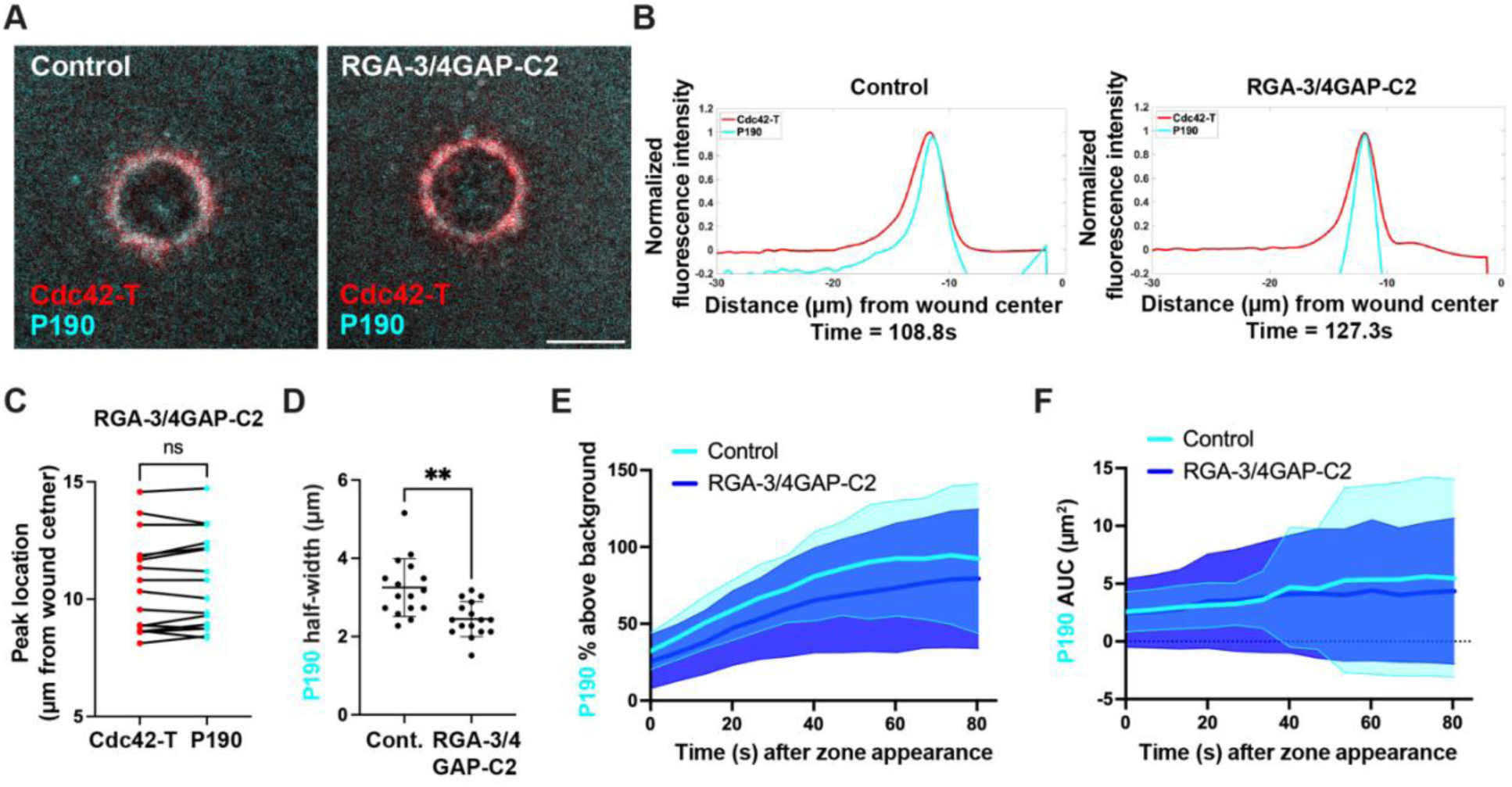
p190RhoGAP polarization to the front of the Cdc42-T zone depends on Rho activity. **(A)** (Left) p190 localizes to the front of the Cdc42-T zone in controls. (Right) p190 localizes throughout the Cdc42-T zone when Rho-T is inhibited via RGA-3/4GAP-C2 expression. **(B)** (Left) spatially averaged and normalized signal intensity of Cdc42-T and p190 from the control micrograph in (A). (Right) spatially averaged and normalized signal intensity of Cdc42-T and p190 from the RGA-3/4GAP-C2 micrograph in (A). Wound center is 0 on the x-axes. **(C)** Cdc42-T and p190 peak location at 20% wound closure in cells expressing RGA-3/4GAP-C2 (n=16). **(D)** P190 half-width at 20% wound closure in controls (n=16) vs. cells expressing RGA- 3/4GAP-C2 (n=16). **(E)** P190 intensity over time after zone appearance in controls (n=17) vs. cells expressing RGA-3/4GAP-C2 (n=20). **(F)** P190 area over time after zone appearance in controls (n=17) vs. cells expressing RGA-3/4GAP-C2 (n=20). (D) shows the mean +/− SD. In (E), and (F), the thick line is the average, and the shaded regions above and below are the SD. A paired t-test was used in (C), and an unpaired t-test was used in (D). **p<0.01. Scale bars for micrographs = 20µm.

## Discussion

The results of this study reveal that cell wounding elicits an astonishing degree of cortical patterning within ∼90s of damage. While this is not primarily a methods study, the extent of cortical patterning would have been impossible to characterize without the Sphinctalyzer. The program’s ability to automatically quantify all the signal in and around the wound permitted exploration of the dynamics of the repair process in unprecedented detail. Moreover, because user input is limited to tracing a single zone in a single frame of the movie, and this is subsequently resampled by the program to find the “perfect” zone peak, essentially all the subjectivity is removed from the analysis. Nor is the Sphinctalyzer just a tool that is only useful for single cell wound analysis: it can potentially be used for anything that has a roughly radial axis of symmetry, such as multicellular wounds, the mitotic spindle (when viewed end-on), the cytokinetic apparatus, and the immunosynapse, to name a few examples.

In addition to the usefulness of this computational tool, the utility of RGA-3/4GAP-C2 for suppression of Rho activity at wounds must also be considered. Wound-induced Rho activation can be suppressed by more traditional approaches such as C3 exotransferase or dominant negative Rho [12,24], but these inevitably reduce cortical Rho activity following injection and thus produce a loss of cortical tension. Because the wound response is qualitatively and quantitatively different in cells with reduced tension [47], experiments must be conducted in a narrow window within which Rho activity is suppressed but the cell is still normally responsive to damage. Pharmacological Rho inhibitors can be applied more acutely, but often suffer from nonspecificity [48]. The use of RGA-3/4GAP-C2 side-steps these issues in that the RGA-3/4 GAP domain is specific for Rho both *in vitro* [49] and *in vivo* (this paper) and, by virtue of its specific targeting to wound edges via the C2 domain, acts in a time- and space-resolved fashion.

One of the results arising from automated analysis of the wound response was the discovery that the closure of the Rho-T and Cdc42-T zones is oscillatory. Previous studies revealed that oscillatory behavior is a feature of the supracellular actomyosin purse strings that form during multicellular wound repair in Drosophila embryos [27], as well as the cytokinetic ring in C. elegans zygotes [50]. Thus, oscillatory behavior may be a general feature of both single and multicellular contractile rings. Further, the fact that the Cdc42-T zone oscillates even when actomyosin-based contractility is suppressed via RGA-3/4GAP-C2 expression indicates that the oscillations do not require contractility per se. This finding is consistent with the observation that cortical Rho GTPase waves can be uncoupled from actomyosin-based contractility in other contexts [30,51] and suggests that the oscillatory behavior of actomyosin in contractile structures may be a downstream consequence of Rho GTPase oscillations.

But the most important finding obtained from automated analysis of oocyte repair was the discovery that the primary Rho GTPase patterning event—segregation of the Cdc42-T and Rho-T zones—is subject to an additional level of patterning wherein the Cdc42-T zone undergoes a distinct front-back polarization of its direct and downstream targets: the Toca-1 peak defines the back of the zone, the Arp2/3, cofilin, cortactin, and p190RhoGAP peaks define the front of the zone, and the Cdc42-T, Pak2 and N-WASP peaks define the middle of the zone (Fig 8A). However, even this simple description does not adequately describe the extent of the patterning, in that the different zones arrive at different “steady state” widths: Cdc42-T, N-WASP and Pak2 converge on widths of ∼5 µm, Toca-1 and cofilin on 4 µm, and Arp2/3 and cortactin on 3 µm (Fig 8A).

**Fig 8.**
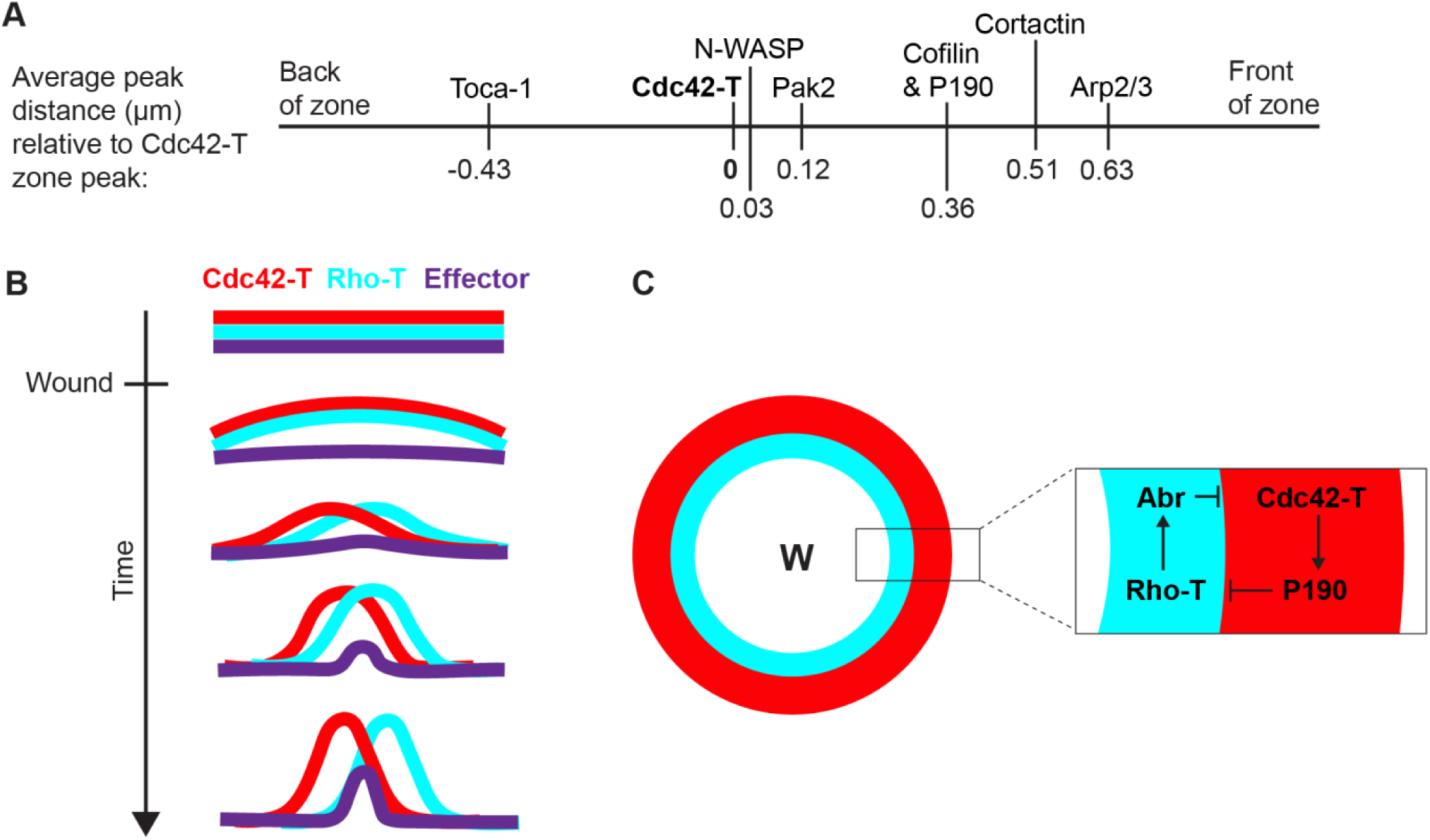
Summary of Cdc42-T effector peak locations and mechanism of Rho GTPase self-organization. **(A)** Comparison of average effector peak distances from the Cdc42-T zone peak based on the position data at 25% wound closure from the main and supplementary figures. Average Cdc42-T peak position was subtracted from the average effector peak position for the sets of experiments in which Cdc42-T was visualized with that effector. This was then repeated for all other groups of experiments in which Cdc42-T was visualized with other effectors. These averages were then plotted relative to the Cdc42-T peak position (0 on the plot). **(B)** Schematic of Rho GTPase self-organization and Cdc42-T effector polarization to the front of the Cdc42-T zone over time. **(C)** Schematic of Cdc42-T and Rho-T mutual exclusion regulated by Abr in the Rho-T zone and p190 in the Cdc42-T zone. This results in self-organization of Cdc42-T and Rho-T into discrete concentric zones around wounds.

How does this polarization occur? Strikingly, the polarized Cdc42-T targets are positioned at boundary regions: Toca-1 concentrates at the boundary between the Cdc42-T zone and the microdomain of PIP2, while Arp2/3, cofilin, cortactin, and p190 concentrate at the boundary between the Cdc42-T zone and the Rho-T zone. This suggests that the combined inputs of the juxtaposed zones are responsible for the observed localization. Consistent with this model, the Toca-1 ΔMGD mutant, which has reduced affinity for Cdc42-T [34] is shifted rearward, away from the Cdc42-T peak and toward the PIP2 peak. Similarly, suppression of Rho activity abolishes the frontward polarization of cortactin and p190RhoGAP.

The recruitment of Cdc42-T direct and indirect targets to boundaries between signaling domains is evocative of the focal expression of specific proteins between developmental compartments in, for example, the Drosophila embryo [52]. While there are many obvious differences relative to cell repair, the important point is that the juxtaposition of two compartments creates a new environment that has a signaling potential that differs from each separate compartment [53]. We thus suggest that boundary development represents a general mechanism for patterning, not only in developing organisms, but also in the cortex of single cells.

A boundary-based mechanism requires the presence of adjacent compartments that differ qualitatively or quantitatively in their constituents, prompting the question of how such compartments arise in the first place. One simple explanation comes from the observation that the Rho-T and Cdc42-T zones are waves, whose wound-ward translocation arises from preferential activation of the GTPases at the leading edge of their zones and preferential inactivation at their trailing edges [30]. Since there must be at least some delay between Rho GTPase activation and resultant activation of their immediate and downstream targets, and since the zones move, spatial differences in signaling protein distribution would arise inevitably [54]. We suspect this explains the relative distributions of PIP2 and Cdc42-T, in that PIP2 begins to accumulate ∼20s after the onset of Cdc42-T accumulation [15].

In contrast, the Rho-T and Cdc42-T zones develop their characteristic juxtaposition via antagonistic self-organization (Fig 8B). As shown here, Cdc42-T indirectly recruits p190RhoGAP to suppress Rho activity in the Cdc42-T zone, while previous work showed that Rho-T recruits the dual GEF-GAP Abr to suppress Cdc42 activity in the Rho-T zone [33] (Fig 8C). Further, with the peculiar circular logic characteristic of self-organization, the localization of p190RhoGAP occurs during polarization of the Cdc42-T zone which, as shown here, is dependent on Rho activity. In other words, the Rho activity at the wound edge is required for the confinement of Rho activity to the wound edge.

How do the current results compare to other models of cell repair? With respect to the temporal patterns of recruitment in the two systems (Dictyostelium and Drosophila) where there is overlap with the proteins studied here, at the temporal level, the Dictyostelium wound response is extremely fast, with N-WASP and Arp2/3 showing consistent recruitment within 2s of wounding and cofilin within 7.5s [18]. In the frog system, the overall temporal pattern of these three proteins is similar, but the response is much slower: 25s for N-WASP, 35s for Arp2/3 and 50s for cofilin. While a different subset of proteins must be compared, the temporal response in syncytial Drosophila embryos is closer to frog: 60s for N-WASP [55] and 45-90s for Rho-T and Cdc42-T [20] in flies vs. 20-40s for Rho-T and Cdc42-T in frogs.

The means by which the concentric Rho-T and Cdc42-T patterning is achieved can also be compared between frog oocytes and Drosophila syncytia: in frog, the patterning arises from a combination of GEFs and GAPs, which arrive at their steady state localization pattern via self-organization ([33] and this study), whereas in Drosophila the patterning is driven largely by GEFs, which appear to be recruited directly to their steady state localization pattern [56]. This difference could explain the more extensive overlap of the Cdc42-T and Rho-T zones in the fly system versus the frog system (compare [12] and [20]) in that in the frog system both local activation and local inhibition are employed.

Lastly, the results leave us with a puzzle: suppression of Cdc42 activity (or expression of a GAP-dead p190) roughly doubles total Rho-T and widens the Rho-T zone, without changing the height (intensity) of the Rho-T zone. In contrast, suppression of Rho activity roughly doubles total Cdc42-T and increases the height of the Cdc42-T zone without changing the width of the Cdc42 zone. Thus, loss of their cognate GAPs makes the Rho zone wider, but the Cdc42 zone taller. The Rho-T zone behavior is easy to explain: without the GAP, Rho-T is free to engage in positive feedback with Abr throughout the so-called playing field—the region around the wound where elevated calcium normally supports both Rho and Cdc42 activity [31,57].

However, the behavior of the Cdc42-T zone is just weird: instead of spreading when Rho activity is suppressed, it simply packs more Cdc42-T into the same total space. Nor can it be argued that Cdc42 activity is somehow limited to a particular region of the playing field, since under different experimental conditions the Cdc42-T zone can form farther from or immediately adjacent to the wound [24]. Rather, it appears that the length-scale control of the Cdc42-T compartment is “zone autonomous”—somehow intrinsic to the Cdc42-T zone itself. How such autonomy is achieved is unknown, but the recent finding that Rho GTPase patterns are stabilized by effector proteins [58] raises the possibility that the length scale of the zone is set by the availability of one or more of its effectors.

## Materials and methods

### DNA and mRNA generation

DNA constructs were generated using the pCS2+ vector as a backbone. Probes for active Rho GTPases (mRFP-wGBD, mCh-wGBD, eGFP-wGBD, dTom-rGBD, and BFP-rGBD) were generated as previously described or based on constructs previously described [12]. eGFP-Pak2, Toca-1-eGFP, eGFP-cofilin, and eGFP-cortactin were generated by cloning full length coding sequences of the respective *Xenopus laevis* proteins into a pCS2-eGFP backbone. 3xeGFP-N-WASP, and 3xeGFP-p190RhoGAP were generated by cloning the full-length coding sequences of the respective *Xenopus laevis* proteins into a pCS2-3xeGFP backbone. The Toca-1ΔMGD-eGFP mutant was generated by performing site-directed mutagenesis on the Toca-1-eGFP construct to convert MGD at amino acids 383-385 to IST, reducing Toca-1ΔMGD affinity for binding Cdc42-T [36]. The untagged p190^R1278A^ mutant was generated by cloning the full-length *Xenopus laevis* p190 coding sequence into a pCS2 backbone and performing site-directed mutagenesis to convert the catalytic arginine at amino acid 1278 to alanine [45,46]. Chn1GAP-C2 was generated as previously described [24]. RGA-3/4GAP-C2 was generated by cloning the N-terminal region and GAP domain of *Xenopus laevis* RGA-3/4 (aa1-267), followed by a flexible linker (SAGGx5), followed by the C2 domain of *Xenopus laevis* protein kinase C β (aa149-287), into a pCS2 backbone [24,59]. All PCR was performed using PfuUltra II Fusion High-fidelity DNA Polymerase (Agilent, Madison, WI). Capped mRNA was generated from linearized DNA using the mMESSAGE mMACHINE SP6 Transcription Kit (Invitrogen, Waltham, MA). 1μL of GTP was added and reactions were incubated for 3-4 hours to improve transcription efficiency. mRNA was purified using the RNeasy Mini Kit (Qiagen, Hilden, Germany).

### AlexaFluor647-Arp2/3 complex protein purification

Native bovine Arp2/3 complex was purified from fresh bovine thymus by ammonium sulfate precipitation and a series of ion exchange chromatography steps (DEAE, Source Q and Source S) followed by gel filtration over a Superdex 200 as described [60]. The gel filtered complex was directly fluorescently labelled on reactive cysteine residues by the addition of 5-fold molar excess of maleimide-Alexa647 dye conjugate. After incubation on ice for 2 hours, the reaction was quenched by adding 2 mM DTT followed by incubation for 30 min. The reaction mixture was then loaded onto an N-WASP-VCA column (5 mM Tris-Cl pH 8.0, 50 mM NaCl, 0.5 mM MgCl2, 0.1 mM ATP, 2 mM DTT) and gradient-eluted over 100 column volumes (10 mM Tris-Cl pH 8.0, 1 M NaCl, 0.5 mM MgCl2, 0.1 mM ATP, 2 mM DTT). Fractions containing the labelled complex were gel filtered (5 mM Hepes pH 7.5, 50 mM KCl, 0.5 mM EGTA, 0.5 mM MgCl2, 0.1 mM ATP, 0.2 mM TCEP) over a Superose 6 column, concentrated, snap frozen with 20% glycerol and stored at −80 °C.

### Oocyte preparation and microinjection

*Xenopus laevis* adult female ovaries were surgically removed based on a protocol approved by the University of Wisconsin-Madison Institutional Animal Care and Use Committee. Ovaries were cut into pieces and washed with 1x Barth’s solution (88mM NaCl, 1mM KCl, 2.4mM NaHCO_3_, 0.82mM MgSO_4_, 0.33mM Ca(NO_3_)_2_, 0.68mM CaCl_2_, 10mM HEPES, with additional 25ug/mL ampicillin, 6ug/mL tetracycline, and 50ug/mL gentamicin sulfate; pH 7.4). Oocytes were treated with 8mg/mL collagenase type I (Life Technologies, Carlsbad, CA) in 1x Barth’s for 1 hour at 16°C and then washed with and stored in 1x Barth’s at 16°C. Stage VI oocytes were selected, and the surrounding follicle layers were removed using forceps. A PLI-100 Pico-injector (Harvard Apparatus, Holliston, MA) was used to inject mRNA into oocytes with a glass needle calibrated to a 40nL injection volume. Following mRNA injection, oocytes were incubated at 16°C for ∼16 hours.

mRNAs were injected at the following needle concentrations: mRFP-wGBD at 0.167mg/mL, eGFP-wGBD at 0.025mg/mL, mCh-wGBD at 0.025-0.33mg/mL, dTom-rGBD at 0.025-0.33mg/mL, BFP-rGBD at 0.1mg/mL, 3xeGFP-N-WASP at 0.2mg/mL, eGFP-Pak2 at 0.25mg/mL, Toca-1-eGFP at 0.25-0.33mg/mL, Toca-1ΔMGD at 0.25-0.33mg/mL, eGFP-cofilin at 0.1mg/mL, eGFP-cortactin at 0.1-0.05mg/mL, 3xeGFP-P190 at 0.2-0.05mg/mL, P190^R1278A^ at 0.2-0.7mg/mL, RGA-3/4GAP-C2 at 0.05-0.033mg/mL, and Chn1GAP-C2 at 0.125mg/mL. AlexaFluor647-Arp2/3 complex protein was a generous gift from Peter Bieling (King’s College London, London, England) and was injected at a needle concentration of 80nM-1µM 0.5-1hour before imaging. C3 exotransferase (Cytoskeleton, Inc., Denver, CO) was injected at a needle concentration of 0.08mg/ml 2-4 hours before imaging [12].

### Imaging and image processing

Oocytes were mounted between two clean No. 1.5 glass cover slips (Globe Scientific, Mahwah, NJ) sealed to a metal cavity slide using vacuum grease (Dow Corning, Midland, MI). Oocytes were imaged using a Prairie View laser scanning confocal system mounted on a Nikon Eclipse Ti and wounded using a 440nm dye laser pumped by a MicroPoint 337nm nitrogen laser (Andor Technologies, South Windsor, CT). A Plan Apo 60x oil objective (NA 1.4) (Nikon, Tokyo, Japan) and immersion oil type-F (n=1.518) (Olympus, Tokyo, Japan) were used. Prairie View (Bruker, Middleton, WI) imaging software was used to acquire time-lapse movies, which were all captured at 2x zoom. Time-lapse movies were acquired as z stacks consisting of 5 or 6 1µm z planes and then average projected in FIJI for display and quantification.

### Data analysis: The Sphinctalyzer

The Sphinctalyzer program is a custom-built, open-source software (BSD-2 license) with a user-friendly GUI and is open to the public via the Sphinctalyzer GitHub repository (https://github.com/uw-loci/BementLabSphinctalyzer), where source code, standalone packages, installation instructions and an in-depth manual can be found. This computational tool employs spatial averaging to quantify fluorescence signal around wounds or anything else with a radial axis of symmetry. In brief, a MATLAB program was developed that allows the user to import a timelapse movie, select a reference frame (i.e. a specific timepoint in which a fluorescent ring is visible encircling the wound) and then manually trace a reference path around the ring (Fig 1A). This trace then undergoes iterative subsampling based on the LiveWire implementation of Djikstra’s algorithm, to refine the shortest, brightest path (S1A Fig). This is similar to the MEDUSA algorithm [27], which is based on the snakes active contour model [26] and which has been used to investigate ring-like dynamics of cell repair in multicellular Drosophila embryo wounds [27]. Once the brightest path has been identified in the reference frame, the Spinctalyzer moves through the entire movie, redrawing a trace to match the shortest, brightest path in the ring at each timepoint. For each timepoint, the average fluorescence intensity is calculated at equidistant locations around the reference path and the information thereby obtained is used to output, in graphical and numerical form such metrics as zone intensity, width (half-width), area (AUC), and location (peak location relative to wound center) over time (S1A Fig). Additionally, it outputs a velocity measurement by quantifying zone translocation over time as well as instantaneous rates of change for all metrics mentioned.

Normalization as percent above background was used to compare intensity, half-width, AUC, and velocity for all proteins involved in repair. Normalization from 0-1 was used in representative line-scans to show protein zone location relative to the wound center. Time to half peak intensity was determined based on the time after wounding at which the signal reached 50% of the peak signal for a given movie. For all graphs displaying data at one time point for RGA-3/4GAP-C2 and C3 experiments, the data were taken from the time at 20% wound closure to account for wound closure stalling due to Rho-T suppression. For all other graphs in which one time point of a time-lapse movie was used, the data were taken from the time at 25% wound closure. GraphPad Prism was used to plot graphs and run statistical tests on the data generated by the Sphinctalyzer.

## Acknowledgements

We would like to thank Aditri Schar for technical contributions to the Sphinctalyzer GUI.

## Supplemental Figures

**S1. Fig.**
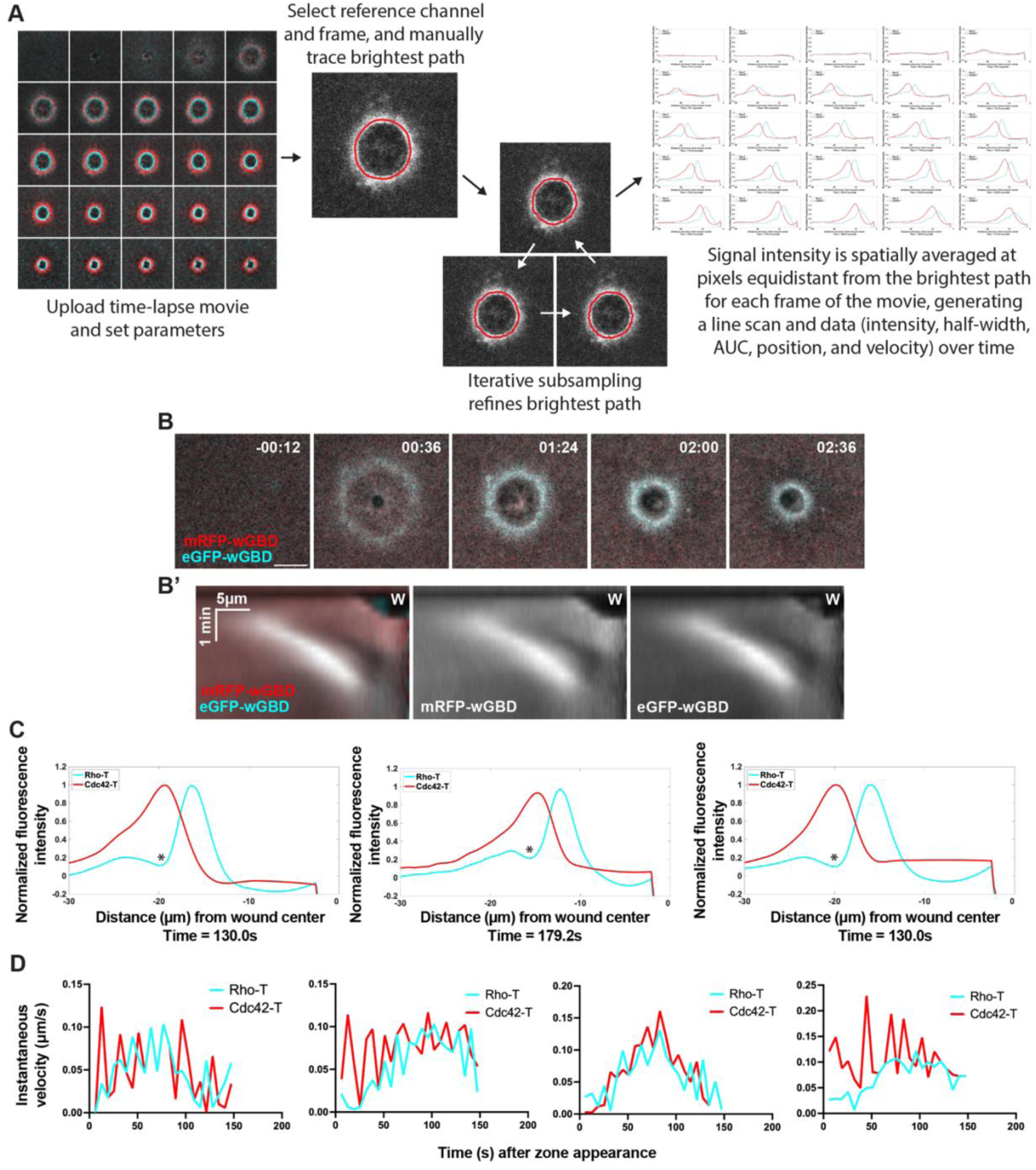
Spatial averaging reveals spatiotemporal dynamics of Rho GTPases during cell repair. **(A)** In-depth overview of Sphinctalyzer workflow and data output. Steps are performed using the Sphinctalyzer GUI (run via MATLAB); see methods and user manual. Time-lapse movie of single cell repair is uploaded to the Sphinctalyzer and parameters such as time interval, microns per pixel, channel names, reference channel, and reference frame are adjusted. The user then draws a trace around the protein zone on the reference image. The Sphinctalyzer uses this trace and performs iterative subsampling based on the Live-wire model to refine the shortest brightest path. The Sphinctalyzer then spatially averages signal intensities at pixels equidistant from the path around the wound for each frame of the movie. This generates a line-scan and intensity, width (half-width), area (AUC), position, and instantaneous velocity measurements for each frame of the movie. **(B)** Montage showing that mRFP-wGBD and eGFP-wGBD dynamically colocalize around wounds over time. Time is in minutes:seconds. **(B’)** Kymograph generated via radial averaging of the movie represented in the montage in (B). W denotes the wound center; only the left half of the radial average is shown. **(C)** Additional examples of the dip in Rho activity in the Cdc42-T zone, denoted by asterisks. Each line-scan is from a different experiment. **(D)** Additional examples of oscillatory changes in Rho-T and Cdc42-T instantaneous velocities over time after zone appearance. Each graph is of one cell from different experiments. Scale bar for micrograph = 20µm.

**S2. Fig.**
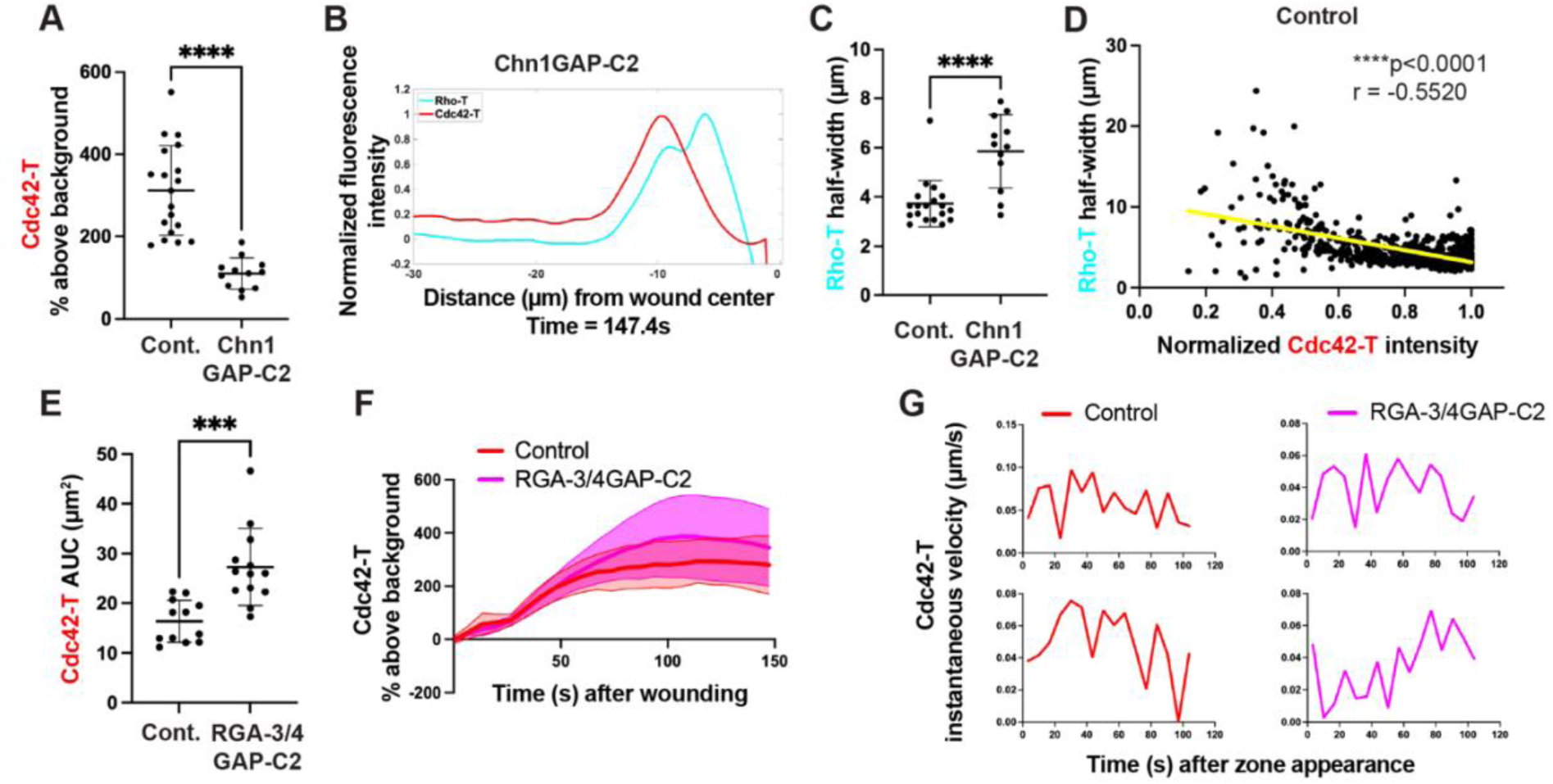
Rho GTPase crosstalk affects Rho-T zone width and Cdc42-T zone translocation. **(A)** Quantification of Cdc42-T zone intensity at 25% wound closure in controls (n=19) vs. cells expressing Chn1GAP-C2 (n=12). **(B)** Additional example of Rho-T zone widening when Cdc42 activity is diminished via Chn1GAP-C2 expression. **(C)** Rho-T zone half-width at 25% wound closure in controls (n=19) vs. cells expressing Chn1GAP-C2 (n=12). **(D)** Correlation of Rho-T zone half-width and normalized Cdc42-T zone intensity in control cells throughout repair (n=32 cells (591 data points)). Cdc42-T zone intensity data are normalized from 0-1to allow for accurate comparison between experiments. **(E)** Cdc42-T zone area at 20% wound closure in controls (n=12) vs. cells expressing RGA-3/4GAP-C2 (n=13). **(F)** Cdc42-T zone intensity over time after wounding in controls (n=13) vs. cells expressing Chn1GAP-C2 (n=13). **(G)** Additional examples of Cdc42-T zone instantaneous velocity oscillations in controls vs. cells expressing RGA-3/4GAP-C2. The top row and bottom row of graphs are from two different experiments. The graphs in (A), (C), and (E) show the mean +/− SD and unpaired t-tests were used. Pearson correlation and simple linear regression were used in (D). In (F), the thick line is the average, and the shaded regions above and below are the SD. ***p<0.001, ****p<0.0001.

**S3. Fig.**
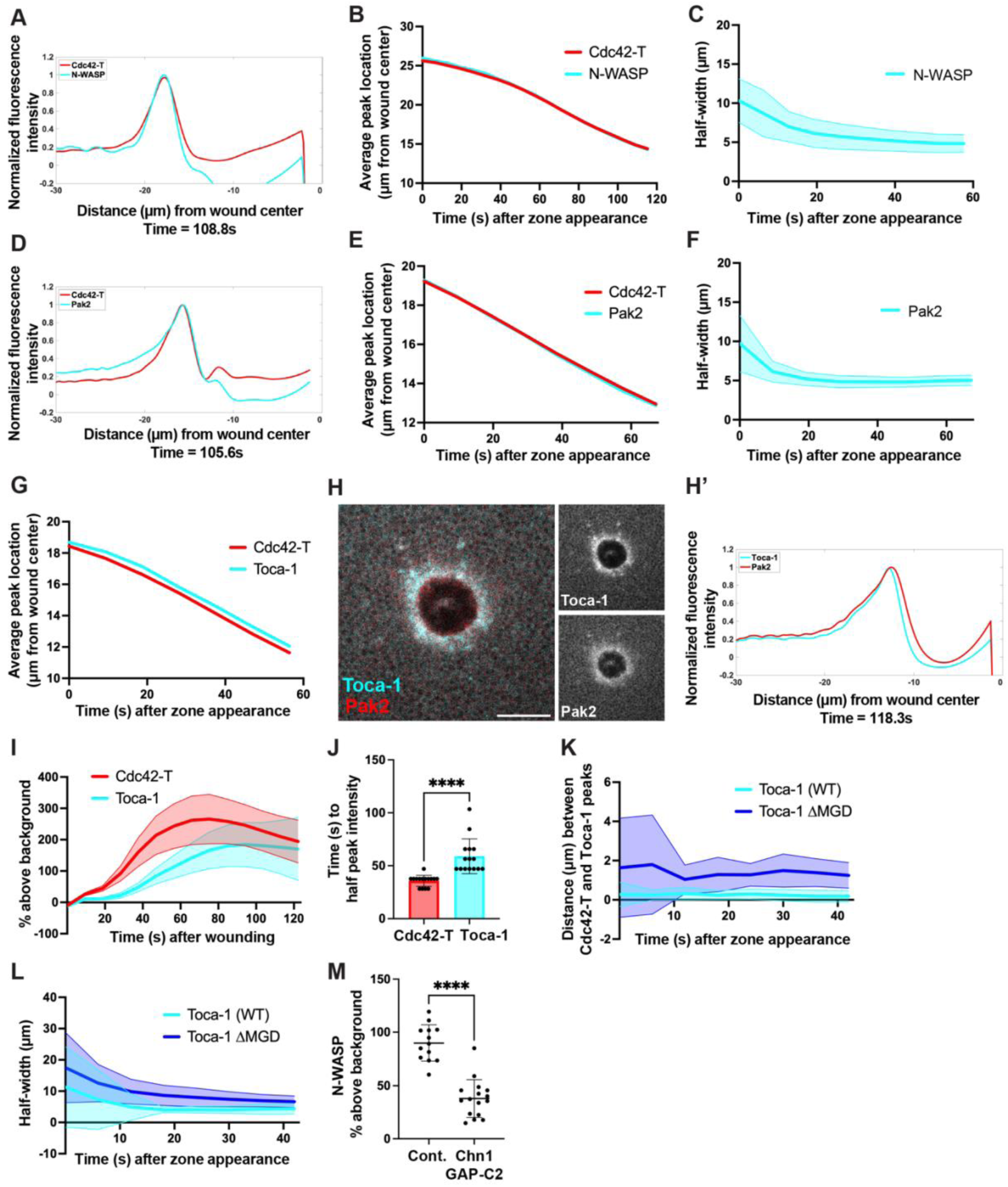
Spatiotemporal dynamics of direct Cdc42-T targets during repair. **(A)** Spatially averaged and normalized signal intensity of Cdc42-T and N-WASP from the micrograph in Fig. 3A. Wound center is 0 on the x-axis. **(B)** Cdc42-T and N-WASP average peak location over time (n=14). **C)** N-WASP half-width over time (n=14). **(D)** Spatially averaged and normalized signal intensity of Cdc42-T and Pak2 from the micrograph in Fig. 3E. Wound center is 0 on the x-axis. **(E)** Cdc42-T and Pak2 average peak location over time (n=15). **(F)** Pak2 half-width over time (n=15). **(G)** Cdc42-T and Toca-1 average peak location over time (n=16). **(H)** Toca-1 and Pak2 enrichment around a wound. **(H’)** Spatially averaged and normalized signal intensity of Toca-1 and Pak2 from the micrograph in (H) showing that Toca-1 polarizes to the back of the Cdc42-T zone even when visualized without a Cdc42-T probe. Wound center is 0 on the x-axis. **(I)** Cdc42-T and Toca-1 intensity over time after wounding (n=15). **(J)** Cdc42-T and Toca-1 time to half peak intensity (n=15). **(K)** Distance between the Cdc42-T and WT Toca-1 peaks (n=18) vs. the distance between the Cdc42-T and Toca-1ΔMGD peaks (n=14) over time. **(L)** Half-width of WT Toca-1 (n=17) vs. Toca-1ΔMGD (n=14) over time. **(M)** N-WASP intensity at 25% wound closure in controls (n=13) vs. cells expressing Chn1GAP-C2 (n=16). The graphs in (J) and (M) show the mean +/− SD. In (C), (F), (I), (K), and (L), the thick line is the average, and the shaded regions above and below are the SD. A paired t-test was used in (J), and an unpaired t-test was used in (M). ****p<0.0001. Scale bar for micrograph = 20µm.

**S4. Fig.**
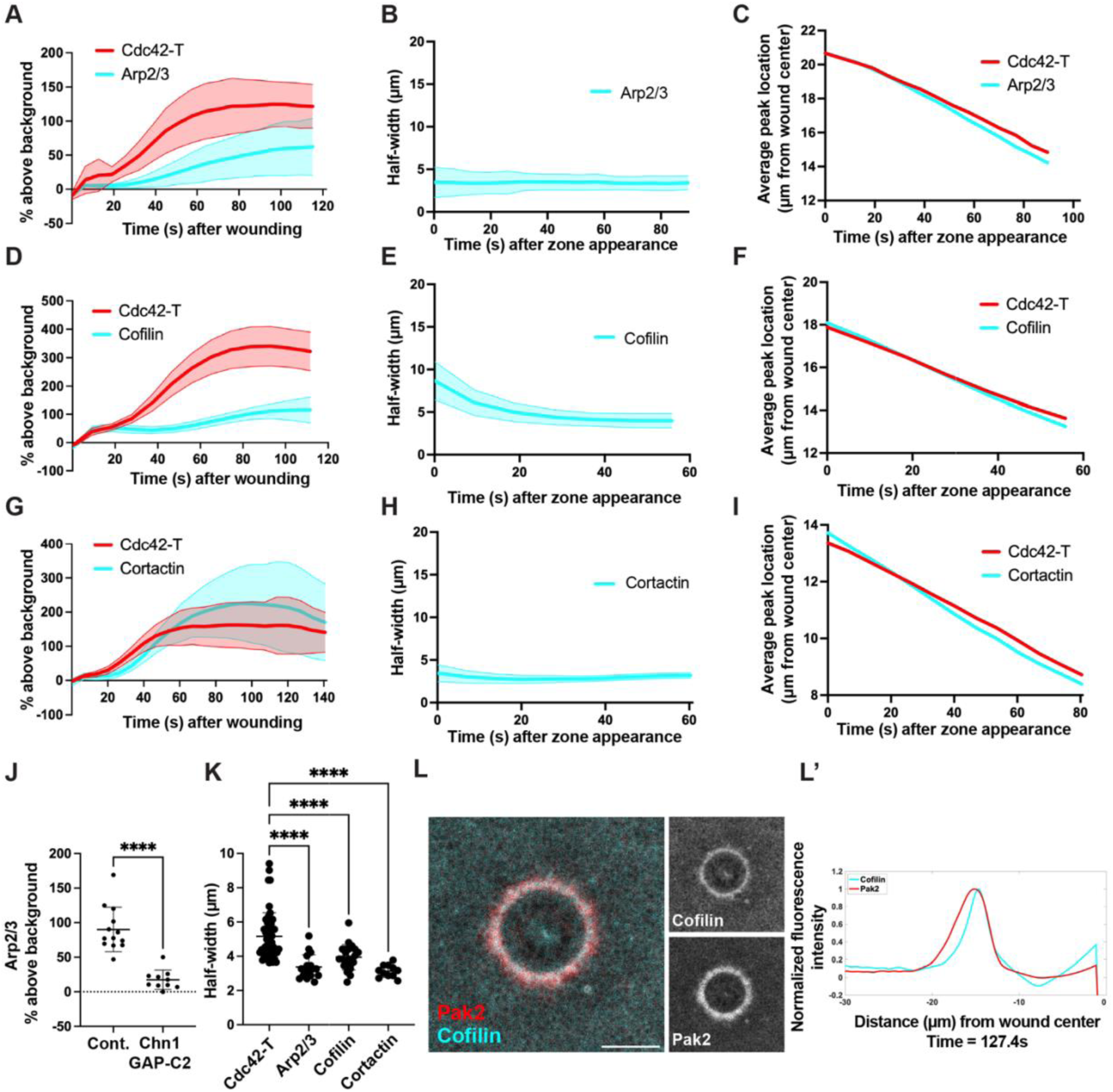
Spatiotemporal dynamics of indirect Cdc42 targets within the polarized Cdc42-T zone. **(A)** Cdc42-T and Arp2/3 intensity over time after wounding (n=17). **(B)** Arp2/3 half-width over time (n=17). **(C)** Cdc42-T and Arp2/3 average peak location over time (n=17). **(D)** Cdc42-T and cofilin intensity over time after wounding (n=22). **(E)** Cofilin half-width over time (n=22). **(F)** Cdc42-T and cofilin average peak location over time (n=22). **(G)** Cdc42-T and cortactin intensity over time after wounding (n=11). **(H)** Cortactin half-width over time (n=11). **(I)** Cdc42-T and cortactin average peak location over time (n=11). **(J)** Arp2/3 intensity at 25% wound closure in controls (n=13) vs. cells expressing Chn1GAP-C2 (n=10). **(K)** Cdc42-T (n=50) vs. Arp2/3 (n=17), cofilin (n=22), and cortactin (n=11) half-widths at 25% wound closure. **(L)** Pak2 and cofilin enrichment around a wound. **(L’)** Spatially averaged and normalized signal intensity of Pak2 and cofilin from the micrograph in (L) showing that cofilin polarizes to the front of the Cdc42-T zone even when visualized without a Cdc42-T probe. Wound center is 0 on the x-axis. (J) and (K) show the mean +/− SD. In (A), (B), (D), (E), (G) and (H), the thick line is the average, and the shaded regions above and below are the SD. An unpaired t-test was used in (J), and a one-way ANOVA with a Dunnett comparison was used in (K). ****p<0.0001. Scale bar for micrograph = 20µm.

**S5. Fig.**
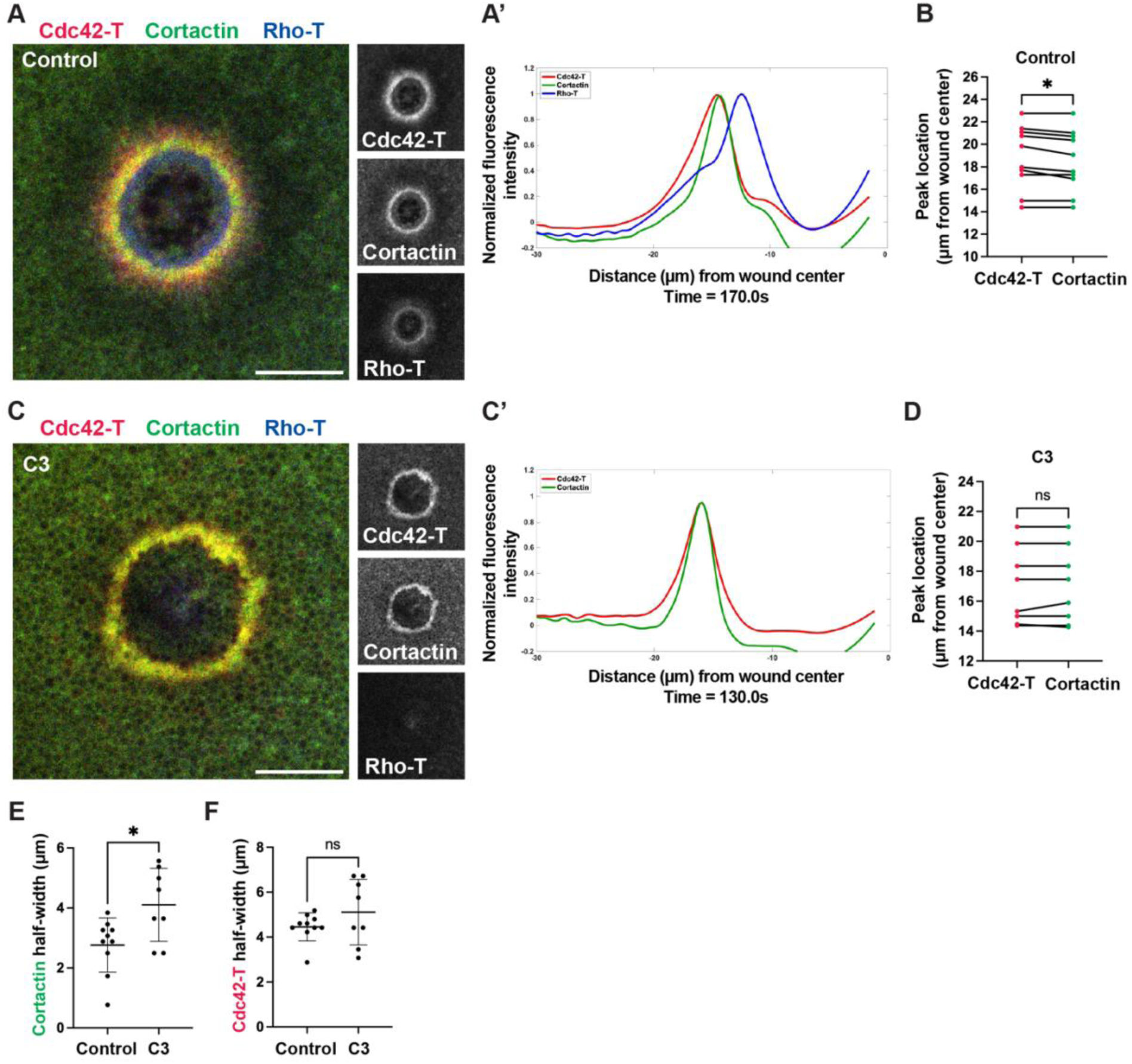
C3-mediated Rho-T suppression recapitulates RGA-3/4GAP-C2-mediated perturbation of Cdc42-T zone polarity. **(A)** Cortactin localizes to the front of the Cdc42-T zone, where it abuts the Rho-T zone. **(A’)** Spatially averaged and normalized signal intensity of Cdc42-T, cortactin, and Rho-T from the micrograph in (A). Wound center is 0 on the x-axis. **(B)** Cdc42-T and cortactin peak location at 25% wound closure in controls (n=10). **(C)** C3 diminishes Rho-T and perturbs Cdc42-T zone polarization, resulting in spread of cortactin throughout the Cdc42-T zone. **(C’)** Spatially averaged and normalized signal intensity of Cdc42-T and cortactin (Rho-T not shown) from the micrograph in (C). Wound center is 0 on the x-axis. **(D)** Cdc42-T and cortactin peak location at 20% wound closure in cells injected with C3 (n=8). **(E)** Cortactin half-width at 20% wound closure in controls (n=10) vs. cells injected with C3 (n=8). **(F)** Cdc42-T half-width at 20% wound closure in controls (n=9) vs. cells injected with C3 (n=8). (E) and (F) show the mean +/− SD. Paired t-tests were used in (B) and (D), and unpaired t-tests were used in (E) and (F). *p<0.05. Scale bars for micrographs = 20µm.

**S6. Fig.**
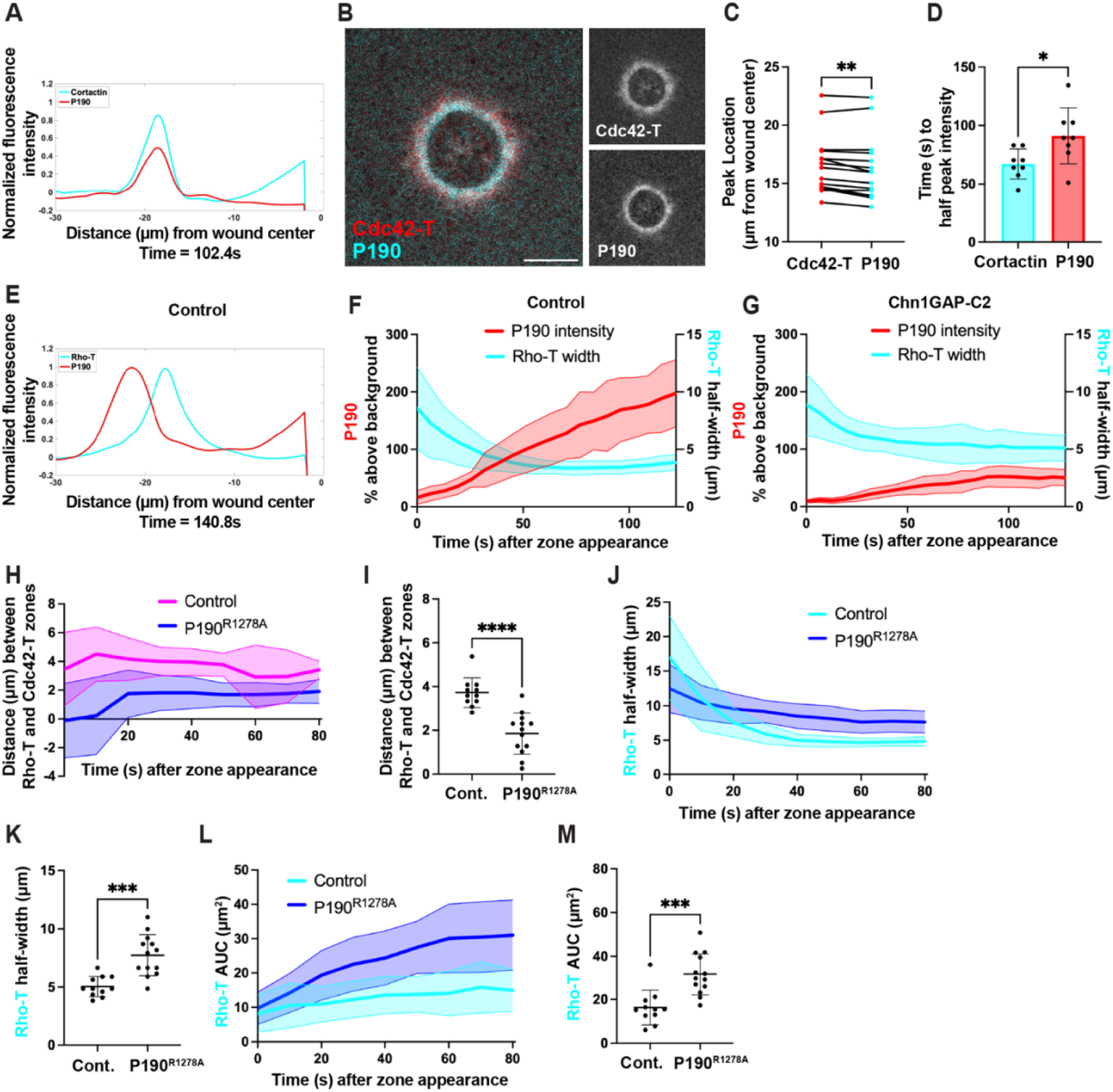
p190RhoGAP polarizes to the front of the Cdc42-T zone thereby confining Rho-T to the wound edge. **(A)** Spatially averaged and normalized signal intensity of cortactin and p190 from the micrograph in Fig 6A. Wound center is 0 on the x-axis. **(B)** P190 localizes to the front of the Cdc42-T zone. **(C)** Cdc42-T and p190 peak location at 25% wound closure (n=15). **(D)** Cortactin and p190 time to half peak intensity (n=8). **(E)** Spatially averaged and normalized signal intensity of Rho-T and p190 from the micrograph in Fig 6D. Wound center is 0 on the x-axis. **(F)** Rho-T zone half-width and p190 intensity over time after zone appearance in controls (n=11). **(G)** Rho-T zone half-width and p190 intensity over time after zone appearance in cells expressing Chn1GAP-C2 (n=8). **(H)** Distance between the Rho-T and Cdc42-T zones over time after zone appearance in control cells (n=11) and cells expressing p190^R1278A^ (n=15). **(I)** Distance between the Rho-T and Cdc42-T zones at 25% wound closure in control cells (n=11) and cells expressing p190^R1278A^ (n=15). **(J)** Rho-T zone half-width over time after zone appearance in control cells (n=11) and cells expressing P190^R1278A^ (n=15). **(K)** Rho-T zone half-width at 25% wound closure in control cells (n=11) and cells expressing P190^R1278A^ (n=15). **(L)** Rho-T zone area over time after zone appearance in control cells (n=11) and cells expressing p190^R1278A^ (n=15). **(M)** Rho-T zone area at 25% wound closure in control cells (n=11) and cells expressing p190^R1278A^ (n=15). (D), (I), (K), and (M) show the mean +/− SD. In (F), (G), (H), (J), and (L), the thick line is the average, and the shaded regions above and below are the SD. Paired t-tests were used in (C) and (D) and unpaired t-tests were used in (I), (K), and (M). *p<0.05, **p<0.01, ***p<0.001, ****p<0.0001. Scale bars for micrograph = 20µm.

## Supplemental Movies

**S1 Movie. Diminishing Rho-T via RGA-3/4GAP-C2 expression results in crawling-like ingression of Cdc42-T zone.** Each time point is an average projection of five opital planes 1µm apart; time points are 6.7s apart. (Left) Rho-T visualized via dTom-rGBD (cyan) and Cdc42-T visualized via eGFP-wGBD (red) zones ingress in coordinated fashion in control cells. (Right) RGA-3/4GAP-C2 expression diminishes Rho-T (cyan) and results in crawling-like ingression of Cdc42-T (red) zone. Elapsed time for control and RGA-3/4GAP-C2 movies is 03:34 (min:s).

**S2 Movie. Toca-1 localizes to the back of the Cdc42-T zone.** Each time point is an average projection of six opital planes 1µm apart; time points are 6s apart. (Left) Toca-1 visualized via Toca-1-eGFP (cyan) concentrates at the back of the Cdc42-T (red) zone visualized via mRFP-wGBD in control cells. (Right) Spatially averaged and normalized signal intensity of Cdc42-T and Toca-1 over time from the micrograph on the left. Wound center is 0 on the x-axis. Elapsed time is 02:24 (min:s).

**S3 Movie. Arp2/3 localizes to the front of the Cdc42-T zone.** Each time point is an average projection of five opital planes 1µm apart; time points are 6.4s apart. (Left) Arp2/3 visualized via AlexaFluor647-Arp2/3 (cyan) concentrates at the front of the Cdc42-T (red) zone visualized via eGFP-wGBD in control cells. (Right) Spatially averaged and normalized signal intensity of Cdc42-T and Arp2/3 over time from the micrograph on the left. Wound center is 0 on the x-axis. Elapsed time is 03:05 (min:s).

**S4 Movie. Cofilin localizes to the front of the Cdc42-T zone.** Each time point is an average projection of six opital planes 1µm apart; time points are 9.3s apart. (Left) Cofilin visualized via eGFP-cofilin (cyan) concentrates at the front of the Cdc42-T (red) zone visualized via mRFP-wGBD in control cells. (Right) Spatially averaged and normalized signal intensity of Cdc42-T and cofilin over time from the micrograph on the left. Wound center is 0 on the x-axis. Elapsed time is 03:06 (min:s).

**S5 Movie. Cortactin localizes to the front of the Cdc42-T zone.** Each time point is an average projection of five opital planes 1µm apart; time points are 6.7s apart. (Left) Cortactin visualized via eGFP-cortactin (cyan) concentrates at the front of the Cdc42-T (red) zone visualized via mCh-wGBD in control cells. (Right) Spatially averaged and normalized signal intensity of Cdc42-T and cortactin over time from the micrograph on the left. Wound center is 0 on the x-axis. Elapsed time is 02:00 (min:s).

**S6 Movie. P190RhoGAP localizes to the front of the Cdc42-T zone.** Each time point is an average projection of five opital planes 1µm apart; time points are 6.4s apart. (Left) P190 visualized via 3xeGFP-p190 (cyan) concentrates at the front of the Cdc42-T (red) zone visualized via mCh-wGBD in control cells. (Right) Spatially averaged and normalized signal intensity of Cdc42-T and p190 over time from the micrograph on the left. Wound center is 0 on the x-axis. Elapsed time is 03:05 (min:s).

**S7 Movie. Cdc42-T and Rho-T localize to mutually exlusive zones.** Each time point is an average projection of six opital planes 1µm apart; time points are 10.3s apart. (Left) Cdc42-T visualized via eGFP-wGBD (red) circumscribes Rho-T visualized via dTom-rGBD (cyan) and Rho-T is confined to the wound edge in control cells. (Right) Spatially averaged and normalized signal intensity of Cdc42-T and Rho-T over time from the micrograph on the left. Wound center is 0 on the x-axis. Note that Cdc42-T and Rho-T occupy two distinct, mutually exclusive zones around the wound. Elapsed time is 02:24 (min:s).

**S8 Movie. P190RhoGAP is necessary for Rho-T zone confinement.** Each time point is an average projection of six opital planes 1µm apart; time points are 10.3s apart. (Left) P190^R1278A^ expression causes Rho-T visualized via dTom-rGBD (cyan) to spread away from the wound edge and into the Cdc42-T (red) zone visualized via eGFP-wGBD, suggesting that p190 is necessary for Rho-T confinement. (Right) Spatially averaged and normalized signal intensity of Cdc42-T and Rho-T over time from the micrograph on the left. Wound center is 0 on the x-axis. Note that Rho-T infiltrates the Cdc42-T zone and mutual exclusion is perturbed. Elapsed time is 02:24 (min:s).

